# Membrane-associated shortened Trk receptors promote neuroprotection and robust axon regeneration without ligands

**DOI:** 10.1101/2022.05.11.491487

**Authors:** Euido Nishijima, Sari Honda, Yuta Kitamura, Kazuhiko Namekata, Atsuko Kimura, Xiaoli Guo, Yuriko Azuchi, Chikako Harada, Akira Murakami, Akira Matsuda, Tadashi Nakano, Luis F. Parada, Takayuki Harada

## Abstract

Activation of neurotrophic factor signaling is a promising therapy for neurodegeneration. However, limited availability of both ligands and receptors permits only transient activation. In this study, we conquered this problem by inventing a new system that forces membrane localization of the intracellular domain of neurotrophin receptor TrkB, which results in constitutive activation without ligands. Our new system overcomes the small size limitation of the genome packaging in adeno-associated virus and allows high expression of the transgene. Single gene therapy using the modified form of TrkB enhances neuroprotection in mose models of glaucoma, and stimulates robust axon regeneration after optic nerve injury. Our system may be also applicable to other trophic factor signaling and lead to a significant advance in the field of gene therapy for neurodegenerative disorders.

## Introduction

Neurodegenerative diseases are debilitating conditions characterized by cognitive and/or motor impairment due to progressive loss of neural function or neuron death. Among them, glaucoma is the leading cause of irreversible blindness due to optic nerve damage and the death of retinal ganglion cells (RGCs). The optic nerve is composed of RGC axons, which transmit visual information from the eyes to brain targets, such as the superior colliculus (SC) and lateral geniculate nucleus. Presently, the reduction of intraocular pressure (IOP) is the sole evidence-based therapy for glaucoma patients, but it is ineffective in a considerable proportion of glaucoma patients, especially those with normal tension glaucoma (1, 2). Thus, other strategies for suppressing further degeneration of RGCs, such as neuroprotection, have been investigated. Gene therapy is potentially an effective therapeutic approach for neurodegenerative diseases, and tools for delivering genes into injured RGCs have been explored (3, 4). Indeed, adeno-associated virus (AAV) delivery of some trophic factors, such as brain-derived neurotrophic factor (BDNF) or ciliary neurotrophic factor (CNTF), stimulates the protection and axon regeneration of RGCs in a mouse model of optic nerve injury (5–7). However, the concentration of such molecules that are exogenously applied is rapidly decreased by diffusion or metabolism, inhibiting sustained signal transduction. Furthermore, the number of receptors available at the cell surface may limit the strength of signal transduction by such trophic factors. Another problem is that the packaging capacity of AAV is limited to ∼4.7 kb, thus some adjustment with the size of promoters and/or molecules will be required to express a large molecule at a high expression level. To overcome these problems, a new idea for effective and continuous stimulation of trophic factor signaling is required. Tropomyosin receptor kinase B (TrkB) is a high-affinity neurotrophin receptor for BDNF that activates several intracellular signaling pathways to promote cell growth and survival upon BDNF binding. Because the extracellular domain of TrkB possesses an autoinhibitory domain (8), we speculate that the membrane-bound intracellular domain of TrkB could induce downstream signaling without BDNF.

Protein lipidation, which includes farnesylation, myristoylation, palmitoylation, and geranylgeranylation, is a method for localizing proteins at the cell membrane by covalent attachment of a lipophilic group. Of those, farnesylation is a form of posttranslational prenylation modification that involves the attachment of a farnesyl group to the C-terminal cysteine residue of the target protein, facilitating membrane association and protein-protein interactions (9, 10). In this study, we generated a constitutive active form of TrkB by farnesylation of the intracellular domain of TrkB (F-iTrkB). The relatively small size of F- iTrkB allowed for the use of the most powerful form of a CAG promoter in an AAV vector, which generated a high expression of F-iTrkB. A single intraocular injection of AAV-F-iTrkB promoted RGC protection and robust axon regeneration without exogenous BDNF application. Our results indicate that artificial lipidation of the intracellular domains of trophic factor receptors triggers powerful signal transduction, which may be effective as a gene therapy tool for neurodegenerative diseases.

## Results

### Membrane anchored intracellular TrkB elicits ligand-independent signaling

We first explored developing constitutively active TrkB. When BDNF binds to TrkB, TrkB dimerizes and transphosphorylates each other, activating downstream signaling pathways, such as Ras-ERK and PI3K-AKT. However, it was previously reported that a TrkB mutant lacking immunoglobulin-domains activated ERK in HEK293 cells without BDNF, but the activity of the TrkB mutant was very low compared with wild-type (WT) TrkB stimulated by BDNF (8). We prepared several TrkB mutants to develop a constitutively active TrkB molecule that induces powerful signal activation (Figure 1A). Analysis of cellular localization revealed that the full-length TrkB (FL-TrkB) was detected at the cell periphery in Neuro2A cells, whereas the intracellular domain of TrkB (iTrkB) and iTrkB with the transmembrane domain (TM- iTrkB) were expressed diffusely in the cytoplasm (Figure 1B). These observations prompted us to design a new TrkB mutant that promotes intracellular membrane anchoring by attaching the farnesylation signal sequence CAAX (F-iTrkB) (Figure 1A). We discovered that F-iTrkB, like FL-TrkB, was localized at the membrane of Neuro2A cells (Figure 1B). We examined the signal transduction activities of these TrkB mutants using immunoblot analysis (Figure 1C). TM-iTrkB and iTrkB failed to activate ERK and AKT, whereas F-iTrkB strongly induced phosphorylation of ERK (pERK) and AKT (pAKT) compared with FL-TrkB alone. Additionally, the levels of pERK and pAKT induced by F-iTrkB were similar to those induced by FL-TrkB stimulated with exogenous BDNF (FL-TrkB+BDNF). Furthermore, F-iTrkB expression increased phosphorylation of other signaling molecules, including Stat1, Stat3, GSK-3β, and p38, as observed by FL-TrkB+BDNF (Figure 1D). We also discovered that myristoylated iTrkB (M-iTrkB) activates ERK and AKT at similar levels to F-iTrkB (Figure 1E).

**Figure 1.**
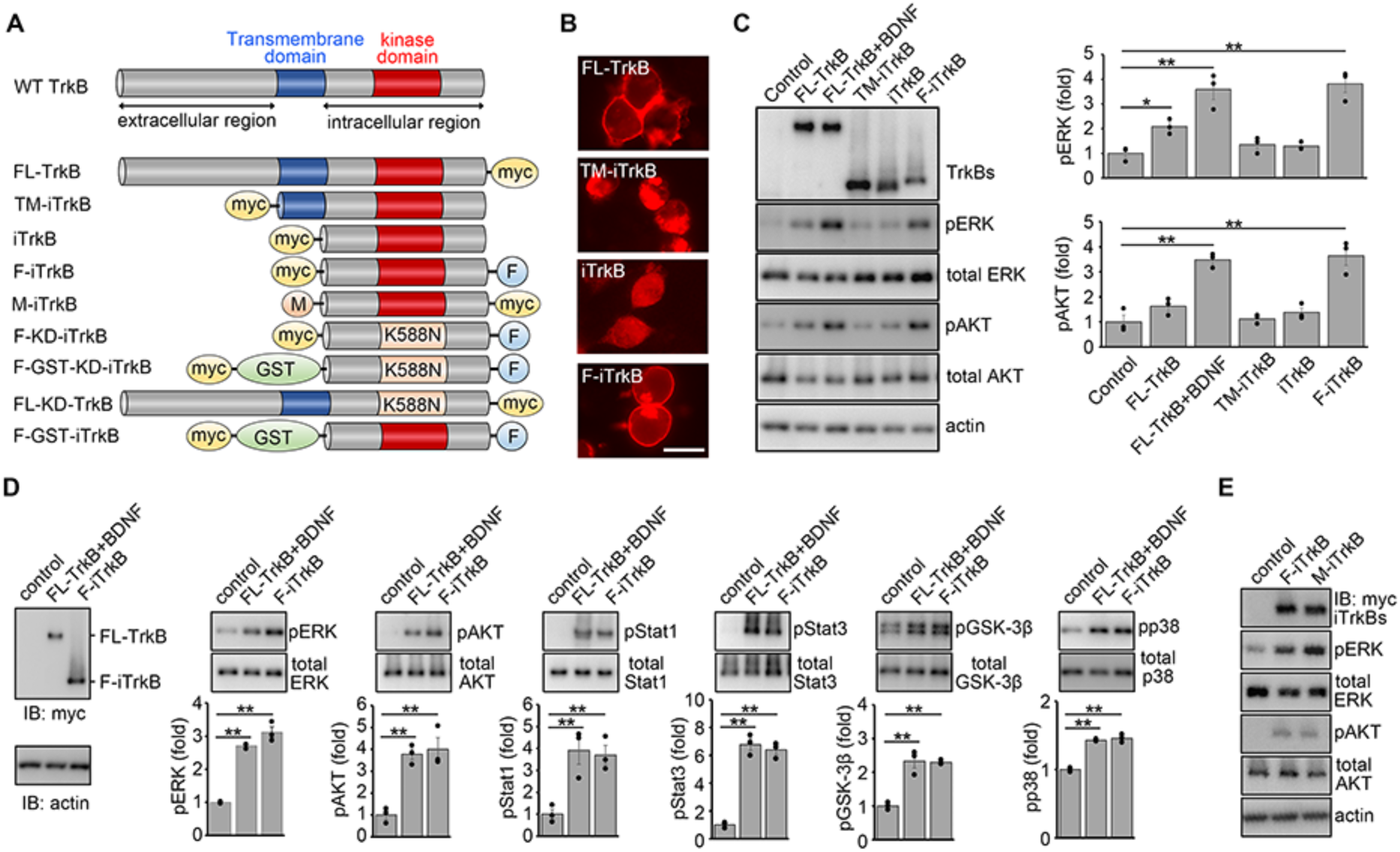
Farnesylated intracellular domain of TrkB activates downstream signaling without ligands. **(A)** Schematic diagram of TrkB constructs used in this study. TM, transmembrane domain; myc, myc-tag; F, farnesylation signal; GST, Glutathione S-transferase; K588N, substitution of lysine with asparagine at position 588, to generate a kinase dead form. **(B)** Cellular localization of TrkB mutants. Immunostaining of myc-tagged TrkB mutants in Neuro2A cells transfected with the indicated plasmids. Full-length (FL)-TrkB and farnesylated intracellular domain of TrkB (F-iTrkB) were localized at the peripheral region. (Scale bar: 25 µm.) **(C)** Activation of ERK and AKT by TrkB mutants. Immunoblot analysis of ERK and AKT phosphorylation in Cos-7 cells transfected with the indicated plasmids. Representative immunoblot images (left) and quantification of the relative levels of phosphorylated ERK (pERK) and AKT (pAKT) (right). The one-way ANOVA with Tukey-Kramer post hoc test was used. *n* = 3 per experimental condition. \*\**P* < 0.01; \**P* < 0.05. **(D)** Comparison of ligand-stimulated FL-TrkB and F-iTrkB in activation of downstream signaling. Immunoblot analysis of several signal proteins in Cos-7 cells transfected with FL-TrkB with BDNF stimulation for 20 min (FL-TrkB+BDNF), or F-iTrkB. TrkBs and actin expression are shown (left). pERK, pAKT, pStat1, pStat3, pGSK-3β and pp38 were detected in both groups. Representative immunoblot images are shown (top), and the relative levels of phosphorylated proteins were quantified (bottom). The one-way ANOVA with Tukey-Kramer post hoc test was used. *n* = 3 per experimental condition. \*\**P* < 0.01. **(E)** The effect of myristoylation and farnesylation on the activity of iTrkB. Cos-7 cells were transfected with myristoylated iTrkB (M-iTrkB) or F-iTrkB. pERK and pAKT were detected from both mutants.

Because F-iTrkB demonstrated powerful activation of multiple downstream signaling without BDNF, we further elucidated its properties. First, we investigated whether the kinase activity of TrkB is essential for F-iTrkB-mediated ERK and AKT activation. Immunoblot analysis revealed that a kinase-dead mutant (F-KD-iTrkB) failed to phosphorylate ERK and AKT, indicating that the kinase activity is essential for F-iTrkB-mediated signal transduction (Figure 2A). We constructed a GST-fused kinase-dead form of F-iTrkB (F-GST-KD-iTrkB) to investigate whether transphosphorylation between F-iTrkB occurs. We discovered that F-GST- KD-iTrkB was phosphorylated at Tyr515 in the presence of F-iTrkB, but not in its absence (Figure 2B), demonstrating that F-iTrkB transphosphorylated F-GST-KD-iTrkB. Additionally, a full-length kinase-dead TrkB (FL-KD-TrkB) was phosphorylated at Tyr515 by F-iTrkB, suggesting that F-iTrkB phosphorylates endogenous WT TrkB (Figure 2C). These results suggest that F-iTrkB transphosphorylates both endogenous WT TrkB and exogenous F-iTrkB at Tyr515. Because BDNF induces TrkB dimerization for transphosphorylation, we investigated whether F-iTrkB forms a dimer using Cos7 cells. A pull-down assay revealed that F-iTrkB does not bind to F-GST-KD-iTrkB (Figure 2D). These results indicate that F-iTrkB does not form a dimer and transphosphorylation of F-iTrkB at Tyr515 was induced by a transient interaction. We also examined whether phosphorylated F-iTrkB is associated with GRB2, Shc, and PLC, like WT TrkB. Immunoblot analysis after a pull-down assay revealed that F-GST-iTrkB was phosphorylated at Tyr515 and bound to GRB2, Shc, and PLC, but not F-GST-KD-iTrkB (Figure 2E). These results indicate that F-iTrkB robustly stimulates downstream signaling without BDNF through the conventional BDNF-TrkB pathway without forming a stable dimer.

**Figure 2.**
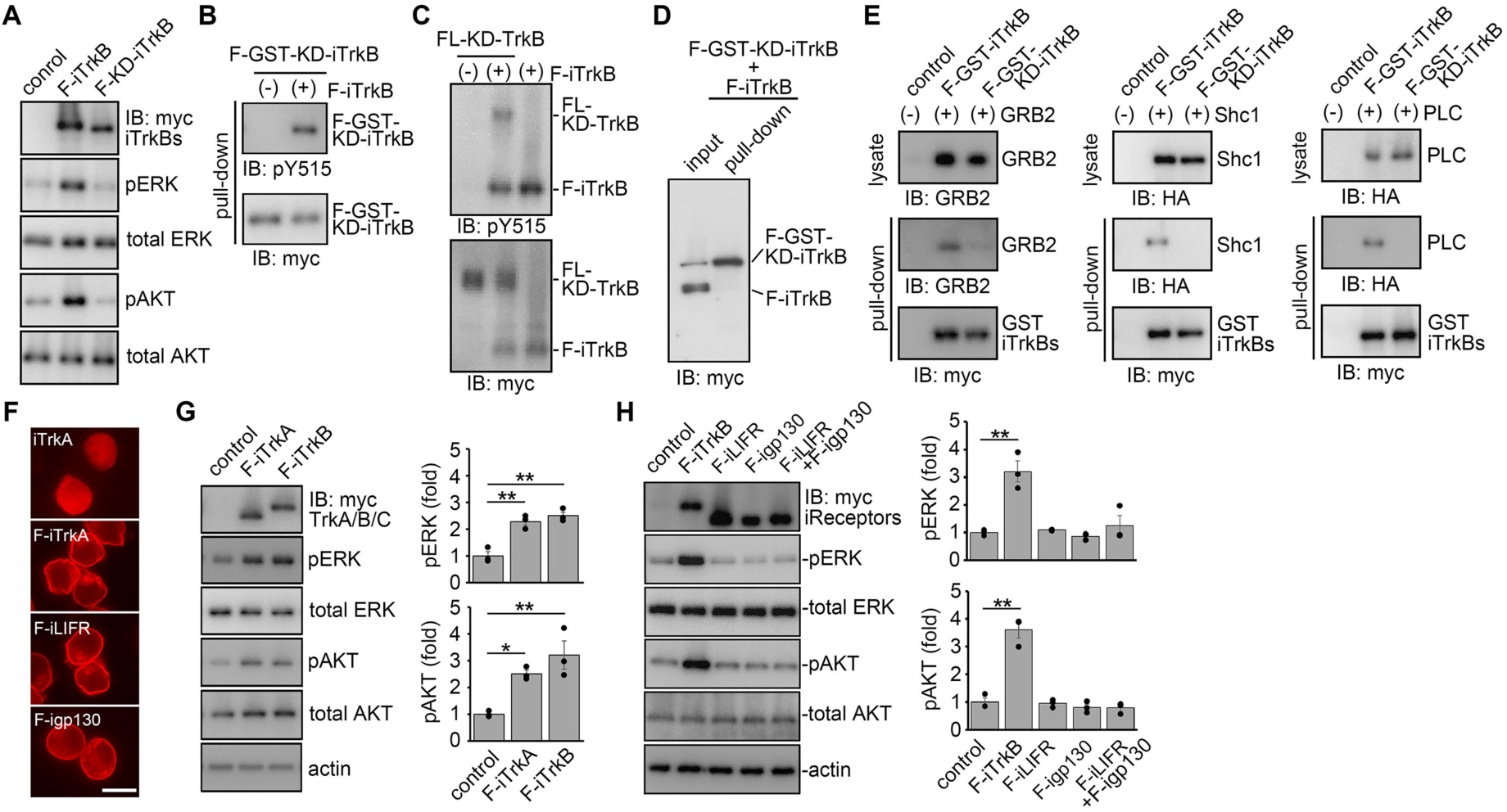
Characterization of F-iTrkB and farnesylated receptors of other trophic factors. **(A)** The effect of kinase activity of F-iTrkB on signal activation. Immunoblot analysis of pERK and pAKT in Cos-7 cells transfected with F-iTrkB or F-KD-iTrkB. Representative images are shown. **(B)** Transphosphorylation of F-iTrkB. Cos-7 cells were cotransfected with a kinase-dead (KD) form of GST-tagged F-iTrkB (F-GST-KD-iTrkB) and F-iTrkB, followed by a GST pull-down assay. The pull-down sample was subjected to immunoblot analysis using an anti-phospho- TrkB (pY515) antibody. **(C)** F-iTrkB-mediated phosphorylation of FL-TrkB. Cos-7 cells were transfected with a KD form of the FL-TrkB (FL-KD-TrkB), alone or cotransfected with F-iTrkB. Total cell lysates were analysed by immunoblotting. **(D)** No detection of F-iTrkB dimers. Cos-7 cells were cotransfected with a KD form of GST- tagged F-iTrkB (F-GST-KD-iTrkB) and F-iTrkB, followed by a GST pull-down assay. The pull-down sample was subjected to immunoblot analysis. **(E)** Interaction of F-iTrkB with GRB2, Shc and PLC. Cos-7 cells were cotransfected with F- GST-iTrkB and GRB2 (left), HA-tagged Shc1 (middle), or HA-tagged PLC (right), followed by a GST-pull down assay. The pull-down sample was subjected to immunoblot analysis. **(F)** Cellular localization of iTrkA, F-iTrkA, F-igp130 and F-iLIFR. Immunostaining of myc- tagged proteins in Neuro2A cells transfected with the indicated plasmids. Farnesylated proteins were localized at the peripheral region. (Scale bar: 25 µm.) **(G)** F-iTrkA-mediated activation of ERK and AKT. Immunoblot analysis of ERK and AKT phosphorylation in Cos-7 cells transfected with F-iTrkA or F-iTrkB. Representative images (left) and quantification of the relative levels of pERK and pAKT (right). The one-way ANOVA with Tukey-Kramer post hoc test was used. *n* = 3 per experimental condition. \*\**P* < 0.01; \**P* < 0.05. **(H)** Absence of signal activation by farnesylated intracellular domain of cytokine receptors. Immunoblot analysis of ERK and AKT phosphorylation in Cos-7 cells transfected with F- iLIFR and F-igp130 and a mixture of the two plasmids. Representative images (left) and quantification of the relative levels of pERK and pAKT (right). The one-way ANOVA with Tukey-Kramer post hoc test was used. *n* = 3 per experimental condition. \*\**P* < 0.01.

Since TrkA is also involved in the survival of neurons, such as RGCs (11), we examined the effect of F-iTrkA. F-iTrkA transfection in Neuro2A cells resulted in membrane localization, but not iTrkA (Figure 2F), and F-iTrkA activated downstream signaling like F-iTrkB (Figure 2G). Furthermore, we examined cytokine receptors, including gp130 and LIFR. To activate cytokine receptor signaling, including CNTF signaling, gp130 and LIFR form a complex. Therefore, we generated F-igp130 and F-iLIFR plasmid constructs. We discovered that transfection of F-igp130, F-iLIFR, and the combination of both (F-igp130+F-iLIFR) failed to activate ERK and AKT, which function downstream of CNTF signaling (Figure 2H). Collectively, these data demonstrate that membrane localization of the intracellular region of TrkA and TrkB activates their downstream signaling pathways in the absence of ligands.

### Overexpression of F-iTrkB alters gene expression in retinal ganglion cells

Death of or damage to RGCs is observed in glaucoma, the second leading cause of blindness globally. BDNF-TrkB signaling plays key physiological roles in the protection of neurons, including RGCs, and decreased expression levels of BDNF and TrkB are observed in the optic nerve head tissues from glaucoma patients (12, 13). Indeed, we discovered accelerated glaucomatous RGC and optic nerve degeneration in aged mice that lack TrkB in neurons (TrkB^c-kit^ KO mice) (14) (Figure 3-figure supplement 1). These findings prompted us to investigate the neuroprotective effects of F-iTrkB on RGCs *in vivo*. For this purpose, we prepared an AAV serotype 2-based vector to express F-iTrkB (AAV-F-iTrkB) or GFP (AAV-GFP) as the control (Figure 3A). Two weeks after intraocular injection of AAV-GFP to the WT mice, we detected numerous GFP-positive cells in the retina (Figure 3A). Following the injection of AAV-F-iTrkB into the eye, immunoblot analysis revealed that F-iTrkB expression was detected in the retinal homogenate (Figure 3B). We performed immunohistological analysis to examine the ability of F-iTrkB to phosphorylate ERK and AKT *in vivo*. Two weeks after intravitreal injection of AAV-F-iTrkB, upregulation of pERK and pAKT was observed in cells expressing myc-tagged F-iTrkB (Figure 3C), indicating that F-iTrkB can induce signal transduction *in vivo*.

**Figure 3 with 1 supplement.**
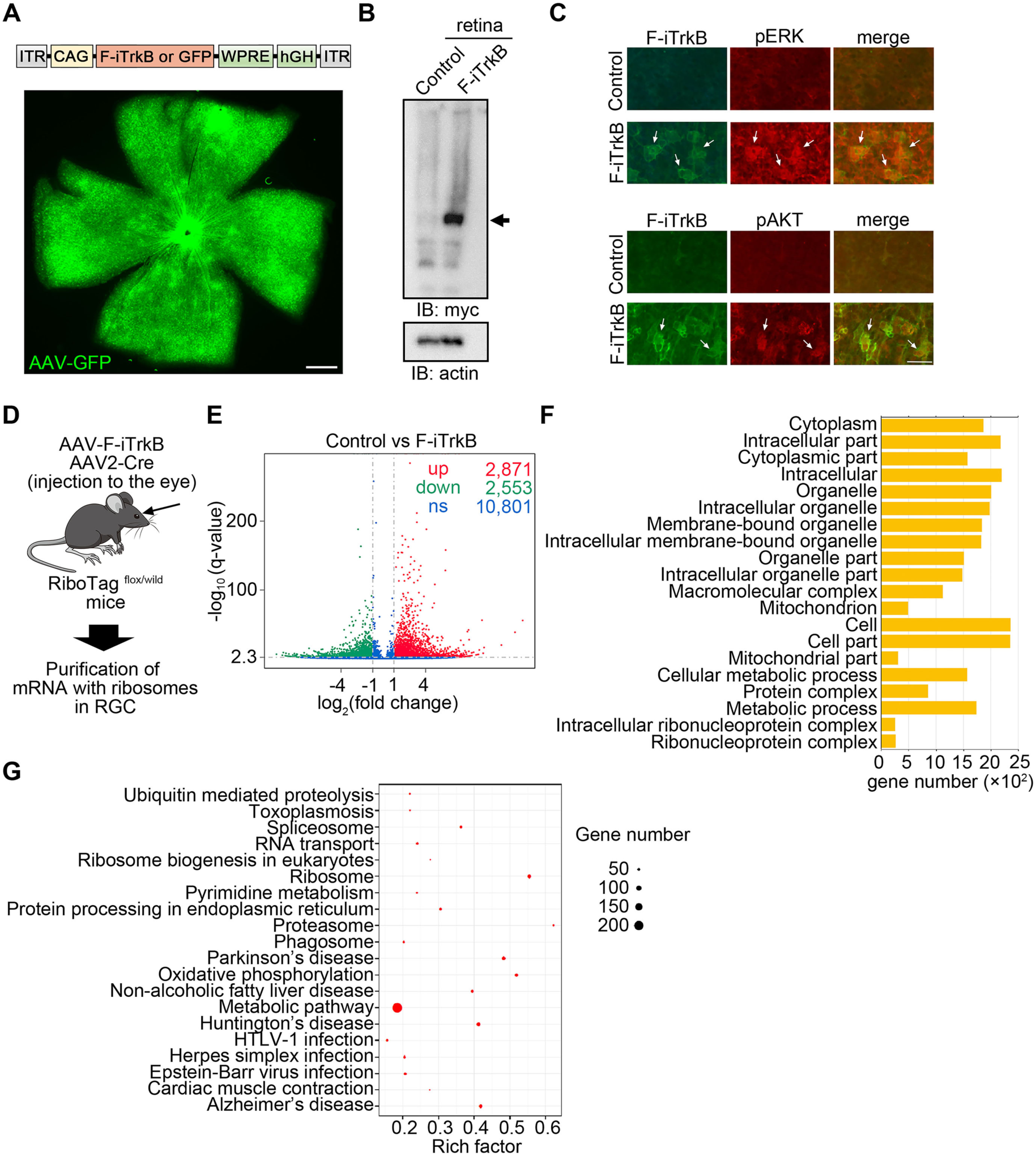
Intravitreal injection of AAV-F-iTrkB alters gene expressions in RGCs. **(A)** Schematic diagram of the plasmid construction for AAV (top). AAV transduction of cells in the mouse retina. AAV-GFP was intravitreally injected to WT mice. Two weeks after injection, GFP expression was detected in the flat-mounted retina (bottom). (Scale bar: 300 µm.) **(B)** Expression of F-iTrkB in the mouse retina. AAV-F-iTrkB was intravitreally injected to WT mice. Two weeks after injection, expression of myc-tagged F-iTrkB were detected in retinal homogenates by immunoblotting. **(C)** F-iTrkB-mediated ERK and AKT activation in RGCs. Double-immunostaining of retinal flat mounts using anti-myc (for F-iTrkB; green) and anti-pERK (top, red) or anti-pAKT (bottom, red) antibodies. (Scale bar: 25 µm.) **(D)** Schematic diagram of the protocol for purification of RGC-specific RNA. **(E)** Volcano plots of gene expression in RGCs infected with AAV-F-iTrkB. The red dots represent significantly upregulated genes, the green dots represent significantly downregulated genes (|log2(Fold Change)| > 1 and q value < 0.005), and the blue dots represent gene expressions with no significant difference between the treatment group (AAV-F-iTrkB) and the control group (AAV-Control). **(F)** Gene Ontology (GO) analysis of differentially expressed genes between AAV-Control or AAV-F-iTrkB-treated RGCs. The most enriched 20 GO terms among the upregulated genes are shown. **(G)** Kyoto Encyclopedia of Genes and Genomes (KEGG) enrichment analysis of differentially expressed genes between AAV-Control and AAV-F-iTrkB-treated RGCs. The top 20 most significantly enriched KEGG pathways are shown.

Next, we investigated the effects of F-iTrkB on gene expressions in RGCs. For this, we isolated mRNAs from RGCs using RiboTag mice (15) (Figure 3D). In RiboTag mice, we intravitreally injected AAV-cre, in combination with AAV-F-iTrkB or AAV-GFP for control. The HA-tagged ribosome was purified by immunoprecipitation using an anti-HA antibody, and subsequently, the ribosome-bound RNA was purified. RNA sequence analysis was performed using purified RNA, revealing that numerous gene expressions were altered by F-iTrkB (Figure 3E). The number of upregulated genes was 2,871, while the number of downregulated genes was 2,553. Among the upregulated genes, the most enriched 20 Gene Ontology (GO) terms (Figure 3F) and the top 20 most significantly enriched Kyoto Encyclopedia of Genes and Genomes (KEGG) pathways are shown (Figure 3G). The data highlighted pathways associated with metabolic processes, such as mitochondrion, mitochondrial part, cellular metabolic process and metabolic process in GO analysis, and oxidative phosphorylation and metabolic pathways in KEGG analysis, implying that F-iTrkB may affect energy homeostasis in RGCs. Furthermore, gene categories associated with protein degradation and neurodegenerative diseases, such as ubiquitin-mediated proteolysis, proteasome, phagosome, Huntington’s disease, and Alzheimer’s disease, were identified in the KEGG analysis. These results suggest that F-iTrkB expression may also affect protein degradation and/or neurodegeneration.

### F-iTrkB protects RGCs in mouse models of glaucoma

To investigate the neuroprotective effects of increased TrkB signaling on RGCs, we first employed a mouse model of normal tension glaucoma, glutamate/aspartate transporter (GLAST) knockout (KO) mice. In GLAST KO mice, RGC degeneration occurs between three and five weeks of age while maintaining normal intraocular pressure (IOP) (16, 17). The relatively fast disease time-course is an advantage when evaluating novel therapies and in this study, we injected AAV-F-iTrkB into the eyeball on postnatal day 10 and examined its effects at three, five, and 12 weeks. IOP was comparable in GLAST KO and WT mice, and intraocular injection of AAV-F-iTrkB did not affect IOP (Figure 4A). We then performed immunostaining of retinal flat mounts using an anti-RBPMS antibody (as a pan RGC maker) and counted the number of RBPMS-positive cells, namely, RGCs (18). AAV-F-iTrkB treatment significantly enhanced the survival of RGCs in 5- and 12-week-old GLAST KO mice (Figure 4B). When we examined retinal morphology *in vivo* using optical coherence tomography (OCT), we discovered that the thickness of the ganglion cell complex (GCC), which contains the RGC layer, was thicker in AAV-F-iTrkB-treated GLAST KO mice than in control mice (Figure 4C). Moreover, visual responses measured by multifocal electroretinography (mfERG) were higher in AAV-F-iTrkB-treated mice than in control mice (Figure 4D). These data indicate that intraocular injection of AAV-F-iTrkB protects RGCs from death and mitigates retinal degeneration and functional decline in GLAST KO mice.

**Figure 4.**
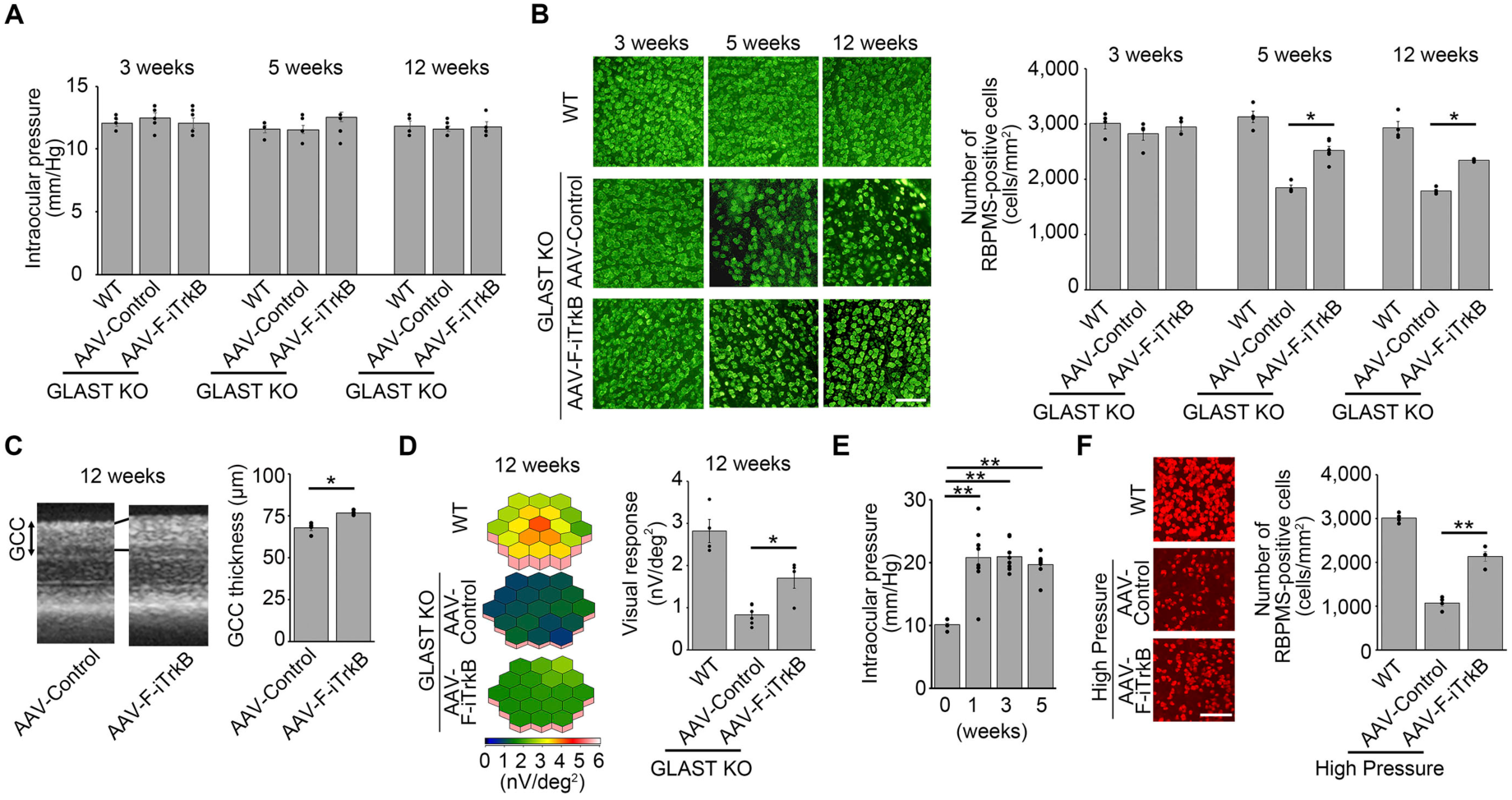
AAV-F-iTrkB prevents RGC degeneration in experimental models of glaucoma. **(A)** Intraocular pressure (IOP) of WT or GLAST KO mice at 3, 5 and 12 weeks old. AAV-F- iTrkB was intravitreally injected in GLAST KO mice at 10 days old. *n* = 4-6 mice per group. **(B)** AAV-F-iTrkB-mediated RGC protection in GLAST KO mice. RGCs were detected by immunostaining of RBPMS in GLAST KO mice. AAV-F-iTrkB was intravitreally injected in GLAST KO mice at 10 days old. Representative images (left) and quantification of RGCs (right). The one-way ANOVA with Tukey-Kramer post hoc test was used. *n* = 4-6 mice per group. \**P* < 0.05. (Scale bar: 100 µm.) **(C)** Optical coherence tomography (OCT) of GLAST KO mouse retinas with or without AAV- F-iTrkB treatment. Cross-sectional images of the retinas in GLAST KO mice at 12 weeks old with or without intravitreal injection of AAV-F-iTrkB (left) and quantification of the ganglion cell complex (GCC) thickness (right). Two-tailed unpaired Student’s *t* test was used. *n* = 4-6 mice per group. \**P* < 0.05. **(D)** Multifocal electroretinography (mfERG) of GLAST KO mice with or without AAV-F- iTrkB treatment. Retinal responses of GLAST KO mice at 12 weeks old with or without intravitreal injection of AAV-F-iTrkB are presented with 3D plots images (left) and quantitative analyses of the retinal response amplitude are shown (right). The one-way ANOVA with Tukey-Kramer post hoc test was used. *n* = 4-6 mice per group. \**P* < 0.05. **(E)** IOP in WT mice with silicone oil-induced ocular hypertension. IOP is elevated from 1 week after the injection of silicone oil into the anterior chamber of the mouse eyes. The one- way ANOVA with Tukey-Kramer post hoc test was used. *n* = 8 mice per group. \*\**P* < 0.01. **(F)** AAV-F-iTrkB-mediated RGC protection in mice with high IOP. RGCs were detected by immunostaining of RBPMS in WT mice at 4 weeks after silicone oil-injection. Representative images (left) and quantification of RGCs (right). The one-way ANOVA with Tukey-Kramer post hoc test was used. *n* = 4 mice per group. \*\**P* < 0.01. (Scale bar: 100 µm.)

Next, we examined the therapeutic effects of AAV-F-iTrkB in a mouse model of high IOP glaucoma. To induce high IOP, we injected silicone oil into the anterior chamber of WT mice to prevent aqueous humor outflow (19). Consistent with the previous report, IOP was elevated chronically from one week after the silicone oil injection (Figure 4E). We injected AAV-F- iTrkB two weeks before the silicone oil injection and analyzed the effects on RGCs four weeks after the induction of high IOP. Compared with a severe RGC loss observed in the control group, AAV-F-iTrkB significantly increased the number of surviving RGCs (Figure 4F). Due to silicone oil interference, we were unable to obtain reliable data from OCT and mfERG. Collectively, these results suggest that AAV-F-iTrkB could be effective in preventing or slowing the progression of glaucoma associated with both high and normal IOP.

### F-iTrkB protects RGCs following optic nerve injury

Furthermore, we examined the effects of AAV-F-iTrkB on RGCs in an acute injury model, the optic nerve crush (ONC) model. ONC was performed two weeks after intravitreal administration of AAV-F-iTrkB, and the number of RGCs was counted in retinal flat mount preparation. Two weeks after ONC, there was a greater number of RGCs in AAV-F-iTrkB- treated mice than in control mice (Figure 5A). These data indicated that AAV-F-iTrkB was also effective in protecting RGCs in an acute retinal degeneration model. Consistent with the reduced number of RGCs, we discovered that synaptic connections, visualized by immunostaining with PSD95 and VGLUT1, also decreased in the control group, but the extent of reduction was milder in AAV-F-iTrkB-treated mice (Figure 5B). The reduced number of synaptic connections could be attributed to cell death and dendritic degeneration. Therefore, we examined the effects of AAV-F-iTrkB on RGC dendritic degeneration. Dendrite morphology was visualized by labeling RGCs with AAV-GFP. We focused on αRGC dendrites rather than dying RGCs because αRGCs are known to be resistant to ONC-mediated cell death (20). RGC dendrites were double-labeled with AAV-GFP and anti-Neurofilament- H (NF-H) antibody, a marker for αRGCs (20) (Figure 5C). Three weeks after ONC, dendrite retraction was observed in αRGCs of the control group, but the extent of retraction was lesser in αRGCs of the AAV-F-iTrkB-treated group (Figure 5D). The total dendritic length, area, and the number of branches significantly decreased after ONC, but AAV-F-iTrkB treatment alleviated these effects (Figure 5E). Sholl analysis revealed that the number of intersections following ONC was significantly higher in AAV-F-iTrkB-treated dendrites than in control dendrites (Figure 5F). These findings indicate that AAV-F-iTrkB treatment minimizes ONC- induced retraction of αRGC dendrites. Accordingly, retinal responses were higher in the AAV- F-iTrkB-treated group than in the AAV-control-treated group after ONC (Figure 5G). These data indicate that AAV-F-iTrkB protects RGC dendrites and synaptic connections after ONC.

**Figure 5.**
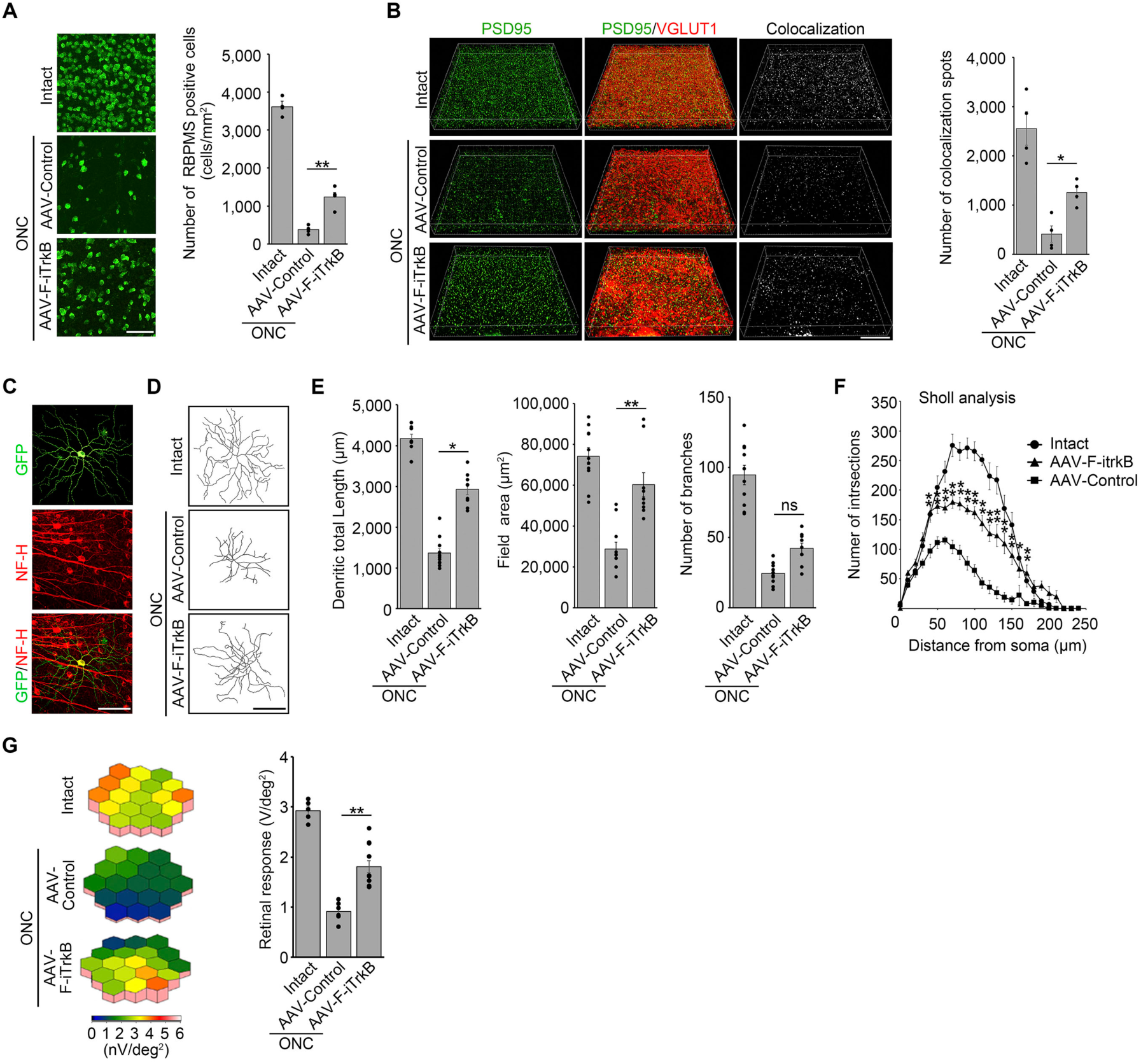
AAV-F-iTrkB prevents RGC degeneration in optic nerve crush injury model. **(A)** F-iTrkB-mediated RGC protection following optic nerve crush (ONC). AAV-F-iTrkB was intravitreally injected in WT mice at 2 weeks before ONC. RGCs were detected by immunostaining of RBPMS at 2 weeks after ONC. Representative images (left) and quantification of the RGC number (right) in intact or injured retinas treated with AAV-Control or AAV-F-iTrkB. The one-way ANOVA with Tukey-Kramer post hoc test was used. *n* = 4 mice per group. \*\**P* < 0.01. (Scale bar: 100 µm.) **(B)** F-iTrkB-mediated synapse protection after ONC. Glutamatergic synapses were visualised in flat-mounted retinas using antibodies against PSD95 and VGLUT1, post- and pre-synaptic markers, respectively. Representative images (left) and quantitative analysis of pre- and post- synaptic co-localized spots (right) in intact or injured retinas treated with AAV-Control or AAV-F-iTrkB. The one-way ANOVA with Tukey-Kramer post hoc test was used. *n* = 4 mice per group. \**P* < 0.05. (Scale bar: 30 µm.) **(C)** Representative images of αRGC morphology. αRGC in the retinal flat mount was double- labeled with AAV-GFP and an anti-Neurofilament-H (NF-H) antibody. (Scale bar: 50 µm.) **(D)** Representative images of αRGC dendritic arbors from intact or injured retinas treated with AAV-Control or AAV-F-iTrkB. (Scale bar: 50 µm.) **(E)** Quantitative analysis of the dendrite length (left), field area (middle), and the number of branches (right) in αRGCs from intact or injured retinas treated with AAV-Control or AAV-F- iTrkB. The one-way ANOVA with Tukey-Kramer post hoc test was used. *n* = 9-11 cells per group. \*\**P* < 0.01; \**P* < 0.05. ns, not statistically significant. **(F)** Sholl analysis of dendrite morphology as a function of distance from the cell soma in intact or injured retinas treated with AAV-Control or AAV-F-iTrkB. The one-way ANOVA with Tukey-Kramer post hoc test was used. *n* = 9-10 cells per group. \*\**P* < 0.01; \**P* < 0.05. **(G)** Retinal responses of mice with intact or injured retinas treated with AAV-Control or AAV- F-iTrkB, measured by mfERG. The 3D plots (left) and quantitative analyses of the retinal response amplitude (right). The one-way ANOVA with Tukey-Kramer post hoc test was used. *n* = 5-12 per group. \*\**P* < 0.01.

### F-iTrkB promotes RGC axon regeneration in an ONC model

Next, we examined the effects of AAV-F-iTrkB on RGC axon regeneration. Two weeks after AAV injection, ONC was performed and Alexa Fluor 647-labeled cholera toxin subunit B (CTB647)-labeled regenerated RGC axons were analyzed. A larger volume of CTB647-labeled regenerated RGC axons was observed in the AAV-F-iTrkB-treated group two weeks after ONC, and this effect was even greater four weeks after ONC (Figure 6A). Four weeks after ONC, some of the regenerated axons reached the optic chiasm, which is approximately 4 mm away from the crush site (Figures 6A and 6B). TrkB activation is widely known to induce downstream signaling pathways, including the PI3K-AKT and Ras-ERK pathways, which are inhibited by PTEN and NF1, respectively. Previous studies have revealed that PTEN deletion induced optic nerve regeneration after ONC (21), but the effects of NF1 deletion are unknown. Therefore, we induced PTEN or NF1 deletion by injecting AAV-cre into the eyes of PTEN^flox/flox^ or NF1^flox/flox^ mice, respectively, two weeks before ONC. We discovered that AAV-F-iTrkB treatment resulted in greater RGC axon regeneration than PTEN deletion (Figures 6A and 6B). These data indicate that F-iTrkB is more powerful than unleashing endogenous levels of PI3K signaling by removing its suppressor PTEN. However, NF1, a Ras- GAP that suppresses Ras signaling, had no effects on RGC axon regeneration (Figures 6A and 6B), indicating that NF1 is not a major Ras-GAP in RGCs.

**Figure 6.**
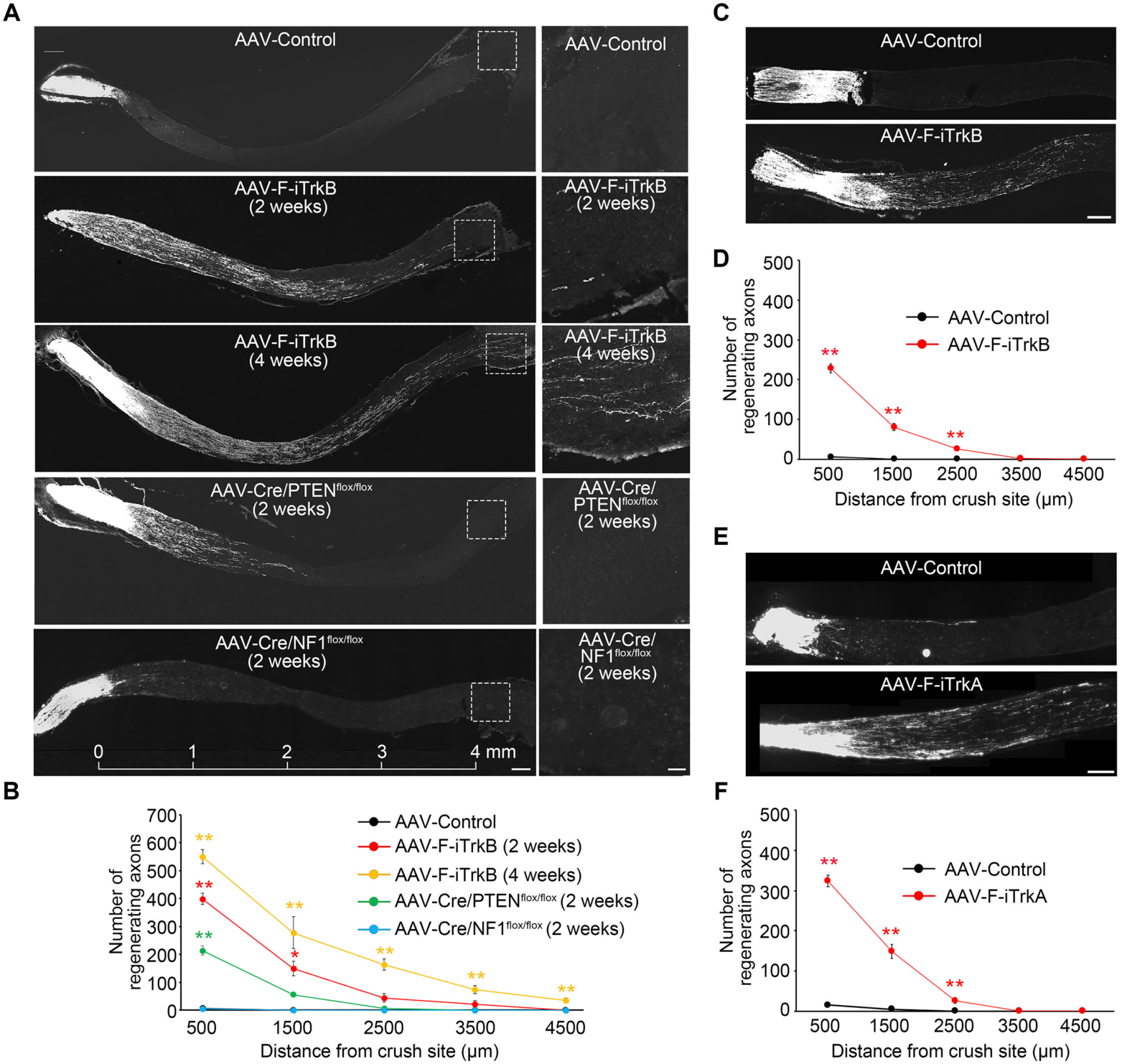
AAV-F-iTrkB promotes robust optic nerve regeneration. **(A)** Representative images of optic nerve sections demonstrating CTB647-labeled regenerating axons in WT mice treated with AAV-F-iTrkB at 2 weeks and 4 weeks after ONC, and in PTEN KO and NF1 KO mice at 2 weeks after ONC (left). Scale bar, 200 µm. Magnification of the boxed areas (right). (Scale bar: 50 µm.) **(B)** Quantification of regenerating axons in the optic nerve shown in (A). The one-way ANOVA with Tukey-Kramer post hoc test was used. *n* = 4 mice per group. \*\**P* < 0.01. **(C)** Representative images of optic nerve sections demonstrating CTB647-labeled regenerating axons in mice that received AAV-F-iTrkB injection intravitreally at 3 min after ONC. (Scale bar: 200 µm.) **(D)** Quantification of regenerating axons in the optic nerve shown in (C). Two-tailed unpaired Student’s *t* test was used. *n* = 4 mice per group. \*\**P* < 0.01. **(E)** Representative images of optic nerve sections demonstrating CTB647-labeled regenerating axons in mice treated with AAV-F-iTrkA at 2 weeks after ONC. (Scale bar: 200 µm.) **(F)** Quantification of regenerating axons in the optic nerve shown in (E). Two-tailed unpaired Student’s *t* test was used. *n* = 4 mice per group. \*\**P* < 0.01.

We injected AAV-F-iTrkB three minutes after ONC to assess its potential for clinical use. AAV-F-iTrkB significantly stimulated the optic nerve regeneration compared with AAV-control, although the effect was not as significant as the pre-administration, and it was similar to PTEN deletion (Figures 6C and 6D). We also examined the ability of AAV-F-iTrkA to stimulate RGC axon regeneration because F-iTrkA strongly activated its downstream signaling pathways *in vitro* (Figure 2G). Intravitreal administration of AAV-F-iTrkA promoted RGC axon regeneration to a similar level as AAV-F-iTrkB (Figures 6E and 6F), indicating that AAV-F-iTrkA is as effective as AAV-F-iTrkB. Collectively, these data demonstrate that AAV-F-iTrkB treatment can protect neurons from disease and injury and mediate robust axon regeneration.

### F-iTrkB promotes RGC axon regeneration in an optic tract transection model

As shown above, F-iTrkB induced robust axon regeneration; however, regenerated axons that reached the optic chiasm were sparse, making it difficult to determine if F-iTrkB can repair the visual pathway effectively after injury. Hence, we employed an optic tract transection model in which the distance between the injury site and the axonal projection site is significantly shorter than in the ONC model. For this, RGC axons in adult mice were cut near the SC (22) (Figure 7A). The CTB647-labeled optic tract was shown in a 3D image via tissue clearing (Figure 7B) and in frozen sections (Figure 7C). In the injured mice, the optokinetic responses (OKR) were lost ten weeks after injury (Figure 7D). Unlike the ONC model, optic tract transection did not induce retinal function loss (Figure 7E), or RGC death (Figure 7F) for at least 12 weeks after injury. AAV-control or AAV-F-iTrkB was injected intravitreally two weeks before optic tract transection, and regenerating axons were visualized with CTB647. No CTB647-labeled axons were detected in the SC of control mice, but CTB647-labeled axons were observed in the SC of AAV-F-iTrkB-treated mice (Figure 7G). We measured OKRs 10– 12 weeks after injury and discovered that optokinetic acuity was slightly higher in the AAV-F-iTrkB-treated group than in the control group (Figure 7H). These data indicate that AAV-F- iTrkB can promote RGC axon regeneration even when the injury site is far from the cell body.

**Figure 7 with 1 supplement.**
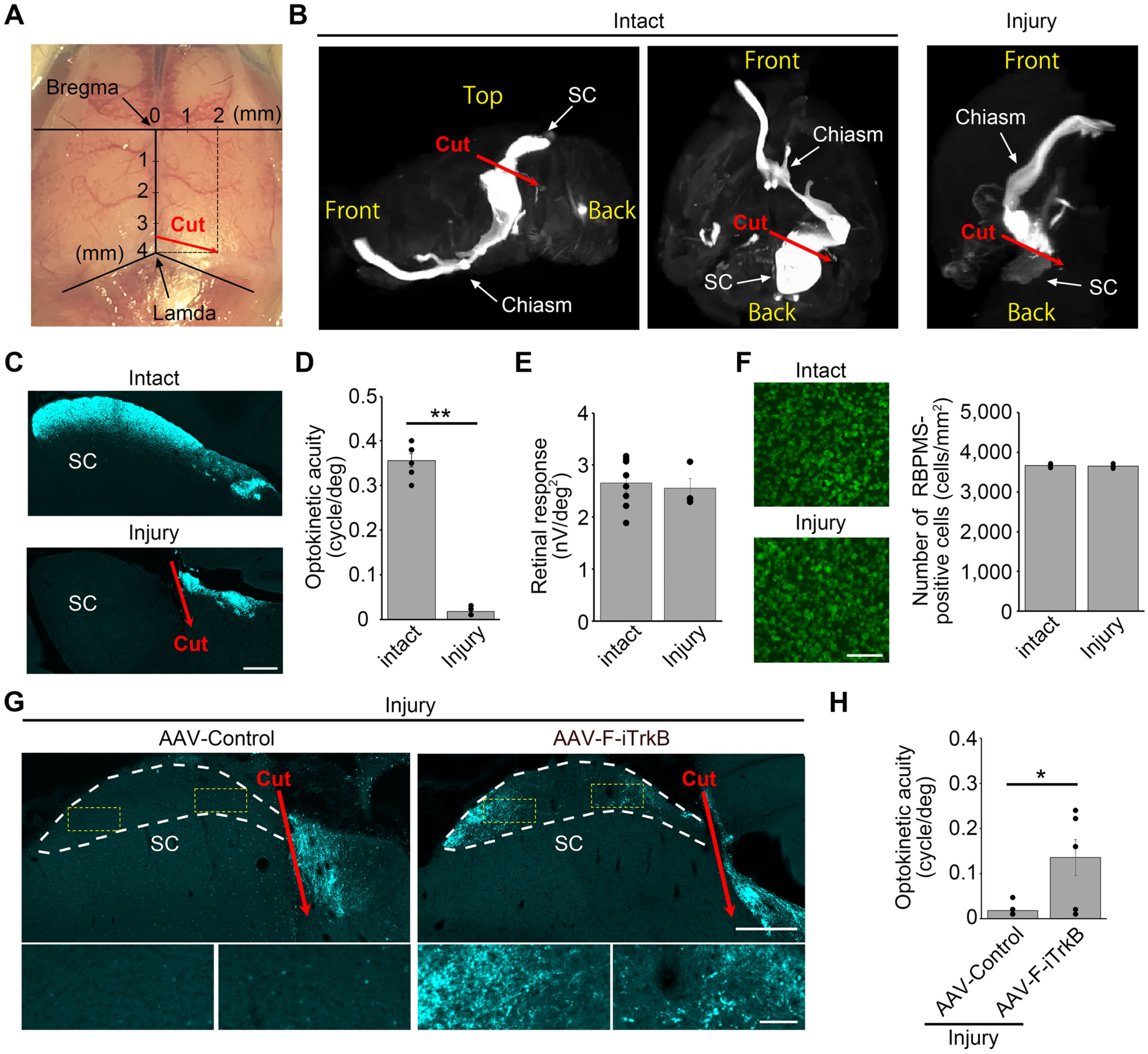
AAV-F-iTrkB promotes axon regeneration in an optic tract transection model. **(A)** A photograph indicating the position of surgical incision for optic tract transection. **(B)** Representative images of the optic tract traced with CTB647 in the intact brain and in the brain that received optic tract transection (injured brain). CTB647 was injected intravitreally and the optic tract from the eye to the superior colliculus (SC) in the brain was made visible by a tissue clearing technique. **(C)** Representative images of sagittal brain sections demonstrating CTB647-labeled axons near the SC in intact and injured brains. (Scale bar: 300 µm.) **(D)** Optokinetic acuity of intact mice or mice with optic tract transection (injured mice). Two- tailed unpaired Student’s *t* test was used. *n* = 4-6 mice per group. \*\**P* < 0.01. **(E)** Averaged retinal responses measured by mfERG in intact and injured mice. *n* = 4-8 mice per group. **(F)** Representative images of retinal flat mounts immunostained with an anti-RBPMS antibody in intact and injured mice at 12 weeks after injury (left). (Scale bar: 100 µm.) Quantification of the number of RBPMS-positive cells (right). *n* = 4-5 mice per group. **(G)** Representative images of sagittal brain sections demonstrating CTB647-labeled regenerating axons near the SC in mice treated with AAV-Control or AAV-F-iTrkB (top). The SC areas were marked with the white dotted lines. (Scale bar: 300 µm.) Magnifications of the boxed areas in yellow are shown at the bottom. (Scale bar: 50 µm.) **(H)** Optokinetic acuity of intact or injured mice treated with AAV-Control or AAV-F-iTrkB. Two-tailed unpaired Student’s *t* test was used. *n* = 6 mice per group. \**P* < 0.05.

## Discussion

In this study, we developed a gene therapy tool, F-iTrkB, which enhances TrkB signaling in neurons in the absence of ligands. Forced localization of iTrkB to the plasma membrane mimicked the activity of FL-TrkB in response to BDNF stimulation. F-iTrkB activates downstream signaling by transphosphorylating each other without forming a stable dimer. The attachment of a farnesyl group to the intracellular domain activated the downstream signaling cascades of TrkB and TrkA. Furthermore, myristoylation, in which a myristoyl group was attached to the N-terminal glycine residue of iTrkB, resulted in a similar significant stimulation of the TrkB signaling pathway. However, the kinase-dead form of F-iTrkB (F-KD-iTrkB) failed to activate ERK and AKT signaling. These data indicate that the orientation of the iTrkB molecule at the plasma membrane has no effect on signal activation, but the kinase activity is essential. The kinase activity is required for F-iTrkB transphosphorylation at Tyr515, causing F-iTrkB to interact with the adaptor proteins Shc and GRB2. This interaction is known to induce multiple complex formations with Ras and PI3K, followed by activation of the Ras-ERK and PI3K-AKT signaling, both of which play crucial roles in neuroprotection and axon regeneration (23–27). Indeed, both F-iTrkB and F-iTrkA successfully activated these signals *in vitro*, and optic nerve regeneration was observed *in vivo* with intraocular injection of AAV-F- iTrkB and AAV-F-iTrkA. These findings imply that the signal transduction cascade of F-iTrkB and F-iTrkA is similar to that of endogenous TrkB and TrkA when stimulated with their ligands. Intraocular AAV-F-iTrkB administration suppressed RGC death in two mouse models of glaucoma and after ONC. Recent studies have revealed that TrkB expression is reduced in aged human and marmoset glaucoma eyes (12, 28). TrkB expression is also reduced in axotomized RGCs in rats, and TrkB gene delivery enhances RGC survival, especially in combination with BDNF application (29). These data indicate that increased TrkB signaling in RGCs could be an effective therapy for glaucoma. However, maintaining a continued supply of BDNF is impractical. In this study, we discovered that AAV-F-iTrkB expression changes numerous gene expressions in RGCs, including genes associated with energy metabolism. Regulation of energy balance is crucial in protecting injured or dying cells and axon regeneration. RGC protection was observed in the high IOP glaucoma model six weeks after AAV-F-iTrkB injection and in GLAST KO mice even after ten weeks, suggesting long-term beneficial effects of a single injection of AAV-F-iTrkB. In addition to the loss of RGC soma, dendrite length and dendritic arbor are reduced in glaucomatous primate RGCs (30, 31). For this, we show that AAV-F-iTrkB protects RGC dendrites from retraction and preserves synapses in a mouse ONC model. These results indicate that AAV-mediated delivery of F-iTrkB could be effective in preventing further RGC death or decreasing the rate of functional decline in glaucoma. Although little is known about the regeneration of mammalian dendrites after injury, recent studies revealed that activating the insulin-dependent PI3K-AKT-mTOR pathway promotes dendrite regeneration in adult mouse RGCs (32). It is interesting to note that F-iTrkB also activates the PI3K-AKT-mTOR pathway, but dendrite regeneration was not apparent after ONC. These findings imply that fine-tuning the correct combination of various signaling pathways is critical. Future studies will focus on the effects of constitutive activation of the insulin receptor signaling on dendrite regeneration and explore the potential of combination therapy with AAV-F-iTrkB.

Adult CNS axons do not usually regenerate after injury, but recent studies indicate that genetic manipulation can put neurons into a regenerative state, implying that CNS axon regeneration is possible (33). Although long-distance RGC axon regeneration accompanied by the recovery of visually guided behavior after ONC has been reported (34, 35), regenerating axons to reach beyond the optic chiasm has been extremely challenging (21, 36, 37). According to some reports, this is due to altered expression levels of guidance cues in the adult-injured CNS, resulting in the misguidance of regenerating axons (38). Literature review indicates that the discovery of the effects of PTEN deletion on axon regeneration was a breakthrough in this field (21), and subsequent studies attempted to achieve greater length and intensity of axon regeneration primarily using combinatory approaches, such as PTEN deletion with manipulation of other genes, including SOCS3 (36), Lin28 (39), and ATF3 (40). We discovered that intraocular injection of AAV-F-iTrkB alone into WT mice induced robust axon regeneration, with some axons reaching optic chiasm four weeks after ONC. This powerful regeneration may be induced because AAV-F-iTrkB could activate Ras-ERK and PI3K-AKT signaling, both of which are involved in promoting axon regeneration. Furthermore, recent studies identified Stat3 as a key molecule in inflammatory stimulation-mediated neuroprotection and axon regeneration in the optic nerve and spinal cord (41, 42). We discovered that F-iTrkB induced powerful activation of Stat3 *in vitro*, implying that F-iTrkB expression may induce an inflammatory stimulated state in RGCs. F-iTrkB-mediated Stat3 signaling may synergistically work with Ras-ERK and PI3K-AKT signaling to promote powerful neuroprotection and axon regeneration. Such a high degree of regeneration by a single gene manipulation is remarkable, and although direct comparison of efficacy across different approaches is difficult due to differences in experimental settings, we believe that the regenerative ability of AAV-F-iTrkB competes with the best available at present.

A reason for the limited regenerative ability of optic nerve axons could be due to variation in RGC subtypes. To date, over 40 RGC subtypes have been identified, and their responses to injury vary (43–45). It is plausible that the RGC subtype with the highest intrinsic potential for axon regeneration has a low chance of survival after ONC or low AAV infectivity. Therefore, additional studies are required to understand this process better and increase the number of regenerating axons by AAV-F-iTrkB. One possible approach is the combination of AAV-F- iTrkB with other strategies. Currently, the long-term delivery of CNTF by an intravitreal implant with encapsulated cells secreting CNTF is in phase II clinical trials for glaucoma (ClinicalTrials.gov Identifier: NCT02862938). CNTF promotes RGC axon regeneration and protects RGCs (6). Since F-igp130 and F-iLIFR failed to activate CNTF receptor signaling, combinatory treatment with a CNTF implant and AAV-F-iTrkB injection could produce synergistic effects that may be effective in a clinical setting. There was a concern that constitutive activity of TrkB could result in tumor growth, but the intravitreal administration of AAV-F-iTrkB did not have abnormal cell growth or negative side effects. With clinical applications in mind, future studies will investigate the use of inducible systems, such as a light-switchable transgene system and a tamoxifen-inducible Cre/loxP system. For long-term observation, we intend to examine the effectiveness of AAV-F-iTrkB gene therapy in marmoset models of ONC and glaucoma (28). The powerful axon regeneration by F-iTrkB encourages the idea that iTrkB may also be effective in treating spinal cord injury. It has been observed that the expression of truncated forms of TrkB (without the intracellular catalytic tyrosine kinase domain) is significantly increased after spinal cord injury, suggesting that this increased expression could limit the availability of BDNF to facilitate axon regeneration (46). Since AAV-F-iTrkB can stimulate intracellular signaling without ligands, F-iTrkB may induce axon regeneration regardless of the increased expression of truncated forms of TrkB.

Conclusively, a single delivery of AAV-F-iTrkB to the retina protected RGCs in animal models of glaucoma and induced robust axon regeneration after axon injury (Figure 7-figure supplement 1). The powerful therapeutic effects achieved with a single dose is a great advantage because it eliminates tissue damage caused by frequent injections. With further characterization and enhancement of delivery, AAV-F-iTrkB may become an effective gene therapy tool for axonal damage and some neurodegenerative diseases, including glaucoma.

## Materials and Methods

### Key resources table

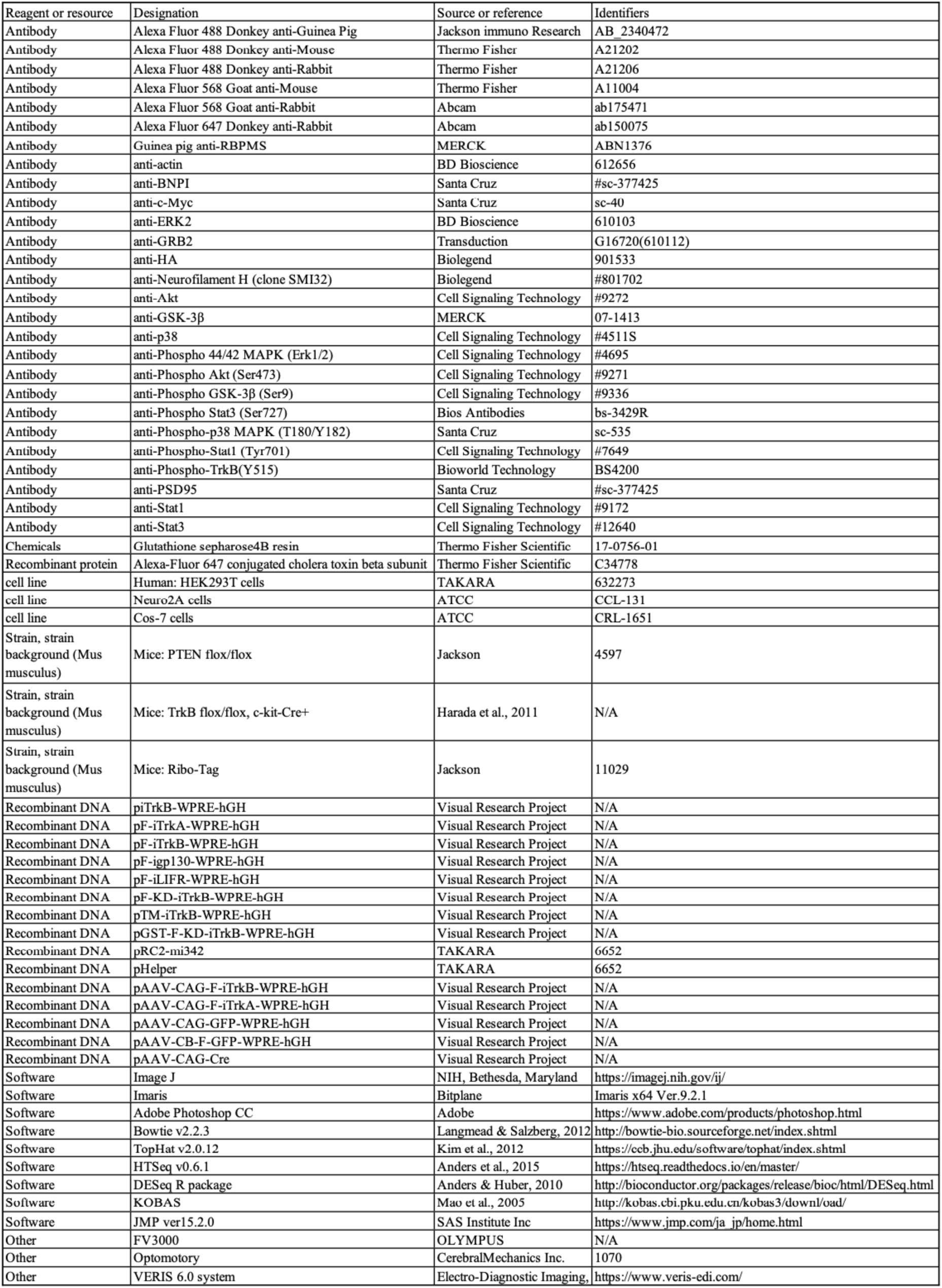

### Animals

Experiments were performed using C57BL/6J, RiboTag (15), TrkB^flox/flox^;c-kit-Cre (TrkB^c-kit^ KO) (14), GLAST KO (17), PTEN^flox/flox^, and NF1^flox/flox^ mice (47). RiboTag and PTEN^flox/flox^ mice were purchased from The Jackson Laboratory. The animals were treated in accordance with the Tokyo Metropolitan Institute of Medical Science Guidelines for the Care and Use of Animals. All animal experiments were approved by the Institutional Animal Care and Use Committee of the Tokyo Metropolitan Institute of Medical Science (18041).

### Plasmids

A plasmid encoding human TrkB was purchased from OriGene Technologies. TrkB mutants were generated by site-directed mutagenesis. The farnesylation-signal of K-Ras or myristoylation signal of Annexin was used for membrane localization (48–50). GRB2, Shc, PLC, TrkA, gp130, and LIFR constructs were obtained from mouse brain cDNA by PCR.

### Transfection and Immunoblot Analysis

Transient transfection in Cos-7 cells or Neuro2A cells was performed using Polyethylenimine HCl Max (Polyscience). After transfection for 20 h, the cells were lysed in SDS-PAGE loading buffer. For signal transduction analysis, cells were serum-starved for 1.5 h before cell lysis. The samples were subjected to immunoblot analysis using antibodies listed in Key resources table. Quantitative analysis was carried out using ImageJ version 2.0.0 (51).

### Pull-Down Assay

Cos-7 cells transfected with plasmids of interest were lysed with a lysis buffer (25 mM Tris pH 7.4, 150 mM NaCl, 1% Triton X-100), and centrifuged at 16,000 × *g* for 10 min. The supernatant was incubated with Glutathione Sepharose 4B resin (Thermo Fisher Scientific) for 30 min at 4°C with gentle agitation. After washing, the precipitated samples were subjected to immunoblot analysis.

### Preparation of AAV

AAVs were produced and purified as described previously (52, 53). Briefly, HEK293 cells were transiently transfected with the AAV vector, pRC2-mi342 and pHelper plasmids (TAKARA). Seventy-two hours after transfection using Polyethylenimine HCl Max, the cells were harvested by scraping, followed by three cycles of freeze-thawing. Cell debris was pelleted with 5,000 × *g* for 20 min, and supernatant were treated with Benzonase (200 U/ ml; Merck) in the presence of 5 mM MgCl_2_ at 37°C for 1 h. The Benzonase-treated viral solution was run on an iodixanol gradient. Purified AAV vectors were washed with Hanks’ balanced salt solution (HBSS) and concentrated using a VIVASPIN 20, 100 kDa MWCO (Sartorius Stedim Lab). Virus titers were determined by quantitative PCR.

### Purification of RGC-Specific RNA

RiboTag mice were injected intravitreally with AAV-Cre for labeling of RGCs. Subsequently, AAV-F-iTrkB were injected intravitreally. Purification of RGC ribosomes was performed as described previously (54, 55), with minor modifications. The mice were perfused transcardially with ice-cold phosphate buffered saline (PBS), and retinas were dissected. Six retinas were pooled as one sample and were homogenized with a Dounce homogenizer in a homogenization buffer (50 mM Tris-HCl, pH 7.5, 100 mM KCl, 12 mM MgCl_2_, 1% Nonidet P-40, 1 mM DTT, 200 U/ml RNAsin, 100 µg/ml cycloheximide, 1 mg/ml heparin). The homogenates were centrifuged at 15,000 × *g* at 4°C for 15 min. The supernatant was incubated with a mouse monoclonal anti-HA antibody (1:50; Biolegend) at 4°C for 16 h. Ribosomes bound to an anti-HA antibody was purified using protein G magnetic beads (GE Healthcare). The RNA bound to ribosomes was purified with RNeasy Protect Mini Kit (QIAGEN) according to the manufacturer’s instructions.

### RNA-Sequencing and Analysis

Sequencing and analysis of the purified RNA was performed by the Novogene NGS Analysis Service (Novogene). Briefly, sequencing libraries were generated using a NEBNext Ultra™ RNA Library Prep Kit for Illumina (New England Biolabs) and index codes were added to attribute sequences to each sample. Clustering of the index-coded samples was performed on a cBot Cluster Generation System using a PE Cluster Kit cBot-HS (Illumina). After cluster generation, the library preparations were sequenced on an Illumina platform and 125-bp/150-bp paired-end reads were generated.

An index of the reference genome was built using Bowtie v2.2.3 (56) and paired-end clean reads were aligned to the reference genome using TopHat v2.0.12 (57). HTSeq v0.6.1 (58) was used to count the read numbers mapped to each gene. Differential expression analysis of two conditions/groups (two biological replicates per condition) was performed using the DESeq R package 1.18.0 (59). *P*-values were adjusted using the Benjamin-Hochberg method. An adjusted *P*-value of 0.005 and log_2_ (fold change) of 1 were set as the threshold for significantly different expression. GO analysis was performed separately for up- and down-regulated gene lists using DAVID Bioinformatics Resources 6.8 (https://david.ncifcrf.gov). KOBAS (60) was used to test the statistical enrichment of differential expressed genes in KEGG pathways.

### Immunostaining of Retinal Flat Mounts and Quantification of RGC Number

Mice were perfused with 4% paraformaldehyde (PFA), and the eyes were enucleated. The removed retinas were first incubated for 2 h in a blocking solution containing with 5% horse serum and 1% Triton X-100 in PBS (pH 7.4). The retinas were then incubated for 24 h with primary antibodies: anti-pERK antibody (1:1000, Cell Signaling), pAKT antibody (1:1000, Cell Signaling) or anti-RBPMS antibody (1:1000, MERCK); followed by incubation with fluorescent-labeled secondary antibodies (Key resources table) at room temperature for 2 h. Images were obtained using the FV3000 confocal microscope (Olympus) or All-in-One fluorescence microscope BZ-X800 (Keyence). The number of RBPMS-positive cells in representative areas (0.04 mm^2^) were counted manually and the average density of RGCs/mm^2^ was calculated (18).

### Retrograde Labeling of RGCs

Retrograde labeling of RGCs was conducted as described previously (61). Briefly, 1% Fluoro-gold (Fluorochrome) dissolved in PBS was injected into the SC using a microsyringe. At 10 days after injection, the mice were sacrificed, and the eyes were enucleated. The retinas were flat-mounted on microscope slides for examination under a BZ-X800 fluorescence microscope (Keyence).

### OCT Imaging

OCT (RS-3000; Nidek) imaging was performed as described previously (17). All line scan images were taken at a distance of three-disc diameters from the optic disc and the average thickness of the GCC (from the inner limiting membrane to the outer boundary of the inner plexiform layer) were measured in retinal images obtained by circular-scanning around the optic disc.

### mfERG

mfERG was performed using a VERIS 6.0 system (Electro-Diagnostic Imaging) as described previously (16, 17). Briefly, the visual stimulus consisted of seven hexagonal areas scaled with eccentricity. The stimulus array was displayed on a high-resolution black-and-white monitor driven at a frame rate of 100 Hz. The second-order kernel was analysed as reported previously (16).

### Induction of IOP Elevation by Intracameral Injection of Silicone Oil

Injection of silicone oil was performed as described previously (19). Briefly, mice were anaesthetized with an intraperitoneal injection of a mixture of medetomidine, midazolam and butorphanol. A 33G needle was tunnelled through the layers of the cornea to reach the anterior chamber without injuring lens or iris. Silicone oil (1,000 mPa.s, Alfa Aesar) was injected slowly into the anterior chamber until the oil droplet expanded to cover most areas of the iris.

### ONC

The mice were anaesthetized with isoflurane during ONC. The optic nerve was exposed intraorbitally and crushed for 5 s at 0.5 mm from the posterior pole of the eyeball with fine surgical forceps (48, 62).

### RGC Dendritic Arbor Imaging

AAV-GFP was administered intravitreally at 2.0 × 10^9^ vector genomes/retina so that the neighbouring RGC dendrites do not overlap with each other. At 2 weeks after AAV-GFP injection, the mice were perfused with 4% PFA, and retinas were immunostained with an anti-NF-H antibody (SMI32 clone, 1:1000; Biolegend) for detection of αRGC. To visualize GFP-labeled dendrites clearly, the immunostained retinal flat mounts were immersed in 85% glycerol (62) and imaged with an FV3000 confocal microscope (Olympus). The obtained *z*-stack images were reconstituted as a 3D images using Imaris software ver 9.2.1 (Bitplane). GFP-labeled dendrites were traced using the Imaris filament tracing function.

### Quantification of RGC Synapses

To visualize RGC dendrites in the inner retinal layer, retinal flat mounts were incubated with anti-BNPI (VGLUT1, 1:1000; Santa Cruz) and anti- PSD95 (1:1000; Cell Signaling) antibodies for 48 h at 4°C with gently agitation, followed by incubation with fluorescent-labeled secondary antibodies (Key resources table). Images were obtained with a FV3000 confocal microscope (Olympus). To reconstruct high resolution 3D images, scans were taken at 0.5-µm intervals with 30 images per focal plane using a 100× objective lens. The number of double-immunolabeled synapses in 3D images were analysed using Imaris.

### Quantification of Regenerating Axons

To visualize regenerating axons in the optic nerve, 2 µl of CTB647 (Thermo Fisher) was injected intravitreally at 2 days before sacrifice. Frozen sections of the optic nerves (14 µm thickness) were obtained by cryosectioning, and CTB647- positive axons was counted manually at 500, 1500, 2500, 3500 and 4500 µm distal to the lesion site. The total number of regenerating axons at different sites in the optic nerve was calculated from the obtained data (48).

### Optic Tract Transection

Optic tract transection was conducted as described previously (22). Briefly, the mice were anaesthetized, shaved, disinfected, and placed in a stereotaxic apparatus. A small incision was made on the scalp and a bone flap was created over the SC. The optic tract was cut with a sharp knife at 3.5 mm posterior to the bregma on the midline to 4 mm posterior to the bregma and 2 mm lateral to the midline at a depth of 2.5 mm (Figure 7A). An antibacterial ointment was applied and the skin was sutured with a 6-0 silk thread.

### 3D Visualization of the Visual Pathway

To visualize the visual pathway in the mouse brain, 2 µl of CTB647 was injected intravitreally. At 2 days after injection, the mice were perfused transcardially with PBS followed by a fixation buffer [4% PFA in 0.1 M phosphate buffer (pH 7.4)]. The whole brain was dissected out and post-fixed with a fixation buffer overnight at 4°C. Tissue clearing was performed by using the 3DISCO method with some modifications (63). Briefly, the brain was incubated in 50% tetrahydrofuran (THF) in distilled water for 12 h, 70% THF for 12 h, 80% THF for 12 h, 100% THF for 3×12 h, and in dibenzyl ether for 2–3 h before imaging. The cleared brain was immersed in dibenzyl ether (refractive index = 1.562) and imaged with a light sheet fluorescence microscope (MVX10-LS; Olympus) (64). We used a 2× objective lens with a 640-nm laser and bandpass filter (660/750 nm). Approximately 500 images were collected by scanning the sample in the *z*-direction with an 8-μm step size. The obtained *z*-stack images were reconstituted as 3D images using Imaris software (ver 9.2.1).

### Visual Behaviour Test

OKRs were analysed using OptoMotry (CerebralMechanics) to measure the highest spatial frequency of the grating tracked, as described previously (22). The mice were placed on an elevated platform surrounded by four computer monitors displaying black-and-white bars. If the head of the mouse moved in concert with the gratings, the trial was scored as ‘tracked’. The optokinetic acuity was determined using an automated staircase procedures. The injured eye refers to the left eye contralateral to the injured right SC. The test normally lasted for 5 min.

### Statistical Analysis

Statistics were performed using JMP 15.2.0 software (SAS Institute). Data are represented as mean ± SEM. Data significance was determined using two-tailed Student’s *t* tests, or one-way ANOVA with Tukey–Kramer post hoc test. Statistical significance is reported as significant at *P* < 0.05.

## Acknowledgements

We would like to thank Jun Horiuchi, Emiko Wakatsuki, Takahiko Noro, Naoki Kiyota, Kaori Segura, Sayaka Ihara, Keiko Okabe, Tomoko Hara, Mayumi Kunitomo, and Yuan Zhu for their technical support and useful discussions. This work was supported in part by Japan Society for the Promotion of Science (JSPS) KAKENHI Grants-in-Aid for Scientific Research (JP21K20979 and JP22K16985 to E.N.; JP20K18404 and JP22K16961 to Y.K.; JP20K07751 to K.N.; JP20K09820 to A.K.; JP21K09688 to X.G.; JP19K09943 and JP22K09804 to C.H.; JP21H04786 to T.H.; JP19KK0229 to T.N. and T.H.; JP21H02819 and JP21K18279 to K.N. and T.H.); and the Takeda Science Foundation, the Suzuken Memorial Foundation, the Naito Foundation, and the Uehara Memorial Foundation (T.H.).

## Author contributions

K.N. and T.H. designed research; E.N., S.H., Y.K., K.N., A.K., X.G., Y.A., and C.H. performed research and analysed data; A. Murakami, A. Matsuda, and T.N. contributed to study discussions and manuscript revisions; L.F.P. contributed new reagents/analytic tools; and E.N., K.N., A.K., L.F.P., and T.H. wrote the paper.

## Data availability

Source data are included in the Source Data files.

---

**Figure 3-figure supplement 1.**
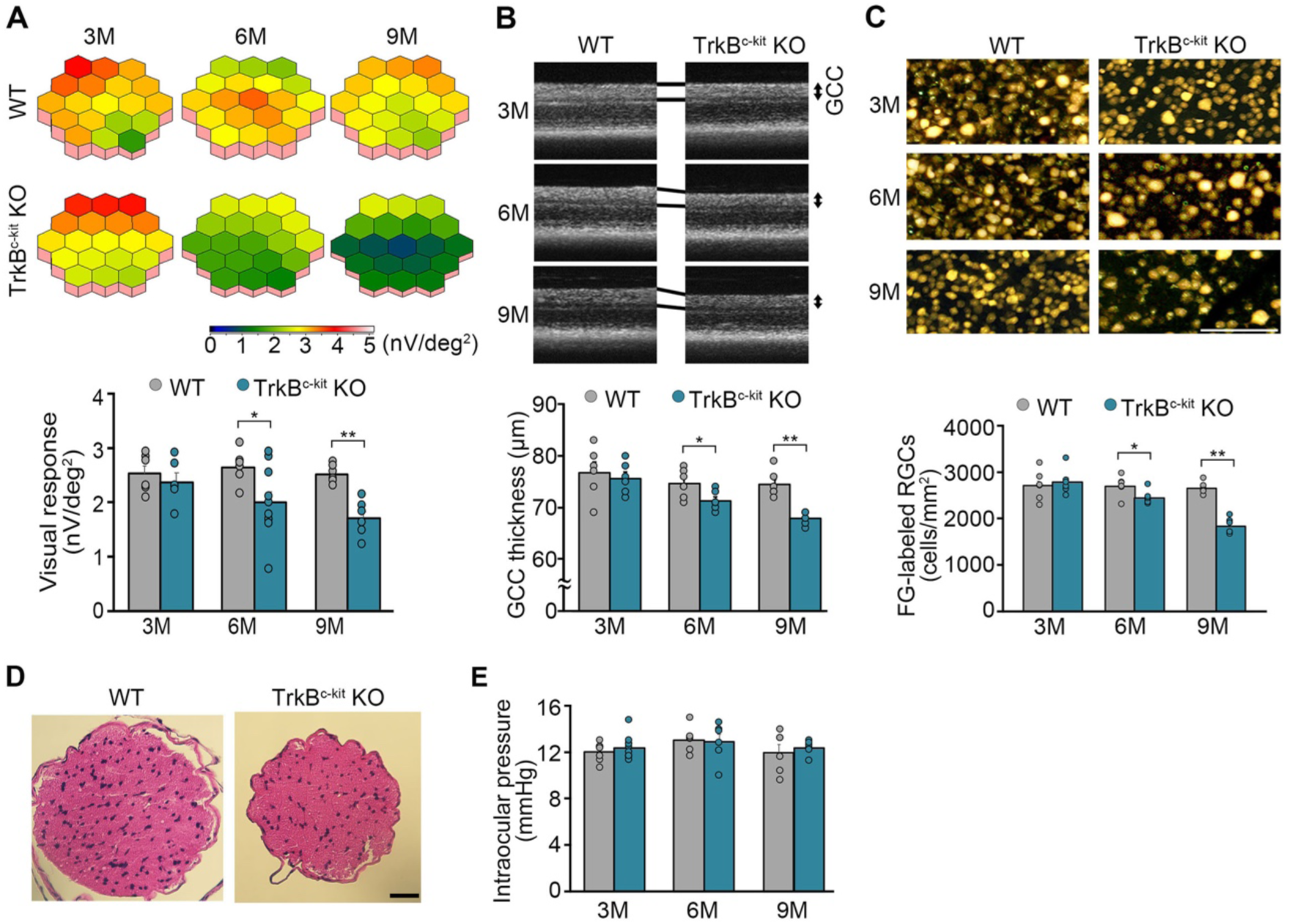
Neuron-specific TrkB deficiency leads to glaucomatous retinal and optic nerve degeneration in aged mice. (A) 3D plots of averaged retinal responses as examined by multifocal electroretinography (mfERG) in WT and TrkB^c-kit^ KO mice at 3, 6, and 9 months (M) of age (upper panel), and quantitative analysis of the visual responses (lower panel). Visual responses in TrkB^c-kit^ KO mice were comparable to those in WT mice at 3 M, but they were significantly reduced by 6 and 9 M compared with WT mice. The one-way ANOVA with Tukey-Kramer post hoc test was used. *n* = 6-10 per group. \*\**P* < 0.01; \**P* < 0.05. (B) Optical coherence tomography (OCT) of WT and TrkB^c-kit^ KO mouse retinas (upper panel), and evaluation of the thickness of the ganglion cell complex (GCC) at 3, 6, and 9 M of age (lower pane). Imaging with OCT revealed that GCC thickness in TrkB^c-kit^ KO mice decreased progressively with time, whereas there was no change in WT mice. The one-way ANOVA with Tukey-Kramer post hoc test was used. *n* = 6 per group. \*\**P* < 0.01; \**P* < 0.05. (C) Retrograde labelling of retinal ganglion cells (RGCs) in WT and TrkB^c-kit^ KO mice (upper panel), and quantitative analysis of Fluorogold (FG)-labelled RGCs (lower panel) at 3, 6, and 9 M of age. RGC number in TrkB^c-kit^ KO mice was significantly decreased compared with WT mice at 6 and 9 M. The one-way ANOVA with Tukey-Kramer post hoc test was used. *n* = 6 per group. \*\**P* < 0.01; \**P* < 0.05. (Scale bar: 100 µm.) (D) Hematoxylin and eosin staining of WT and TrkB^c-kit^ KO mouse optic nerves. Thinning of the optic nerve was observed in TrkB^c-kit^ KO mice at 9 M. (Scale bar: 25 µm.) (E) Intraocular pressure of WT and TrkB^c-kit^ KO mice. Both WT and TrkB^c-kit^ KO mice at 3, 6, and 9 M showed no changes in intraocular pressure with aging. *n* = 6-10 per group.

**Figure 7-figure supplement 1.**
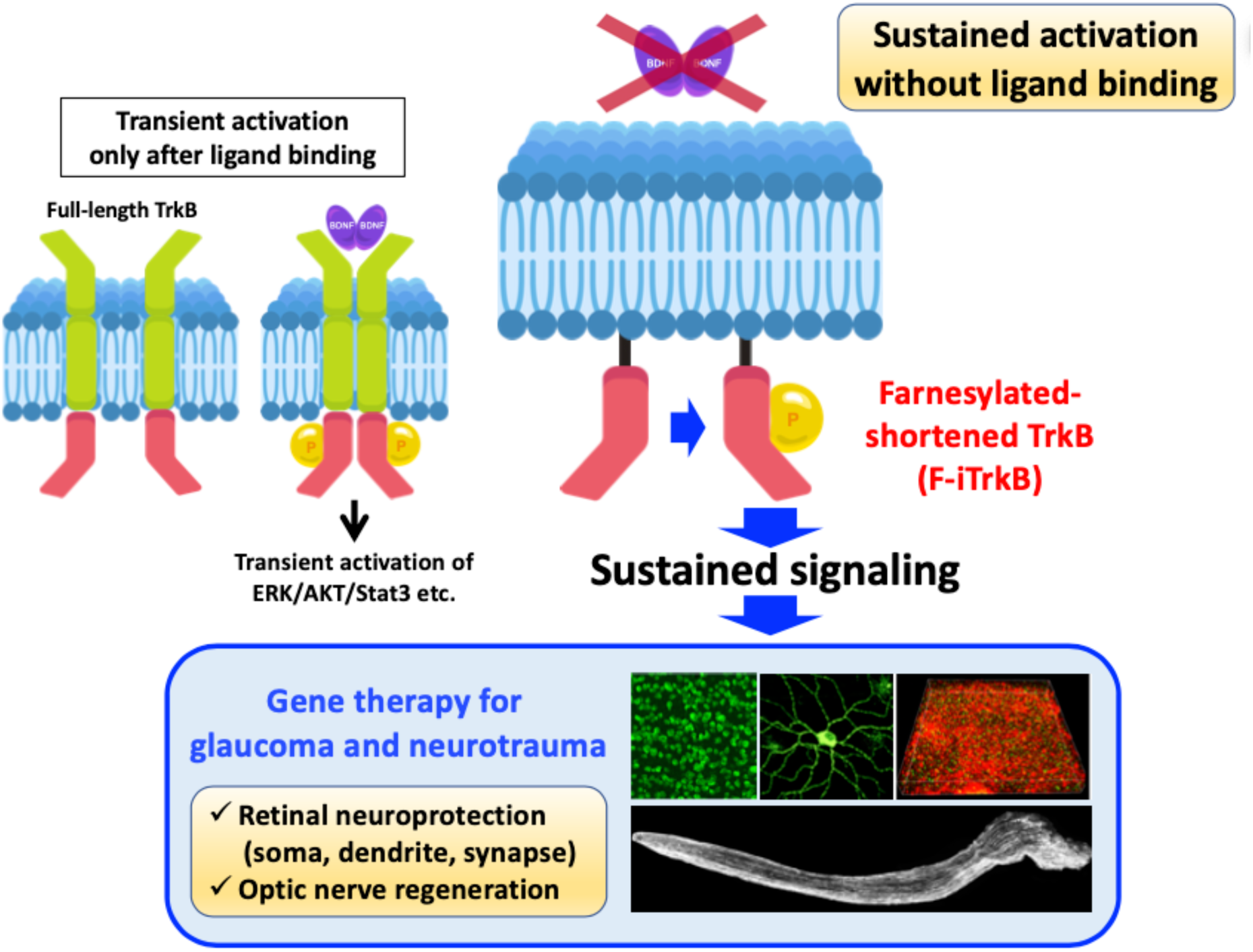
Schematic model of sustained activation of TrkB signaling without BDNF. AAV-mediated delivery of farnecylated-shortened TrkB (F-iTrkB) induced sustained activation of the downstream signaling of full-length TrkB (ERK/AKT/Stat3 etc.) in retinal ganglion cells (RGCs) without BDNF. Intraocular injection of AAV-F-iTrkB induced RGC protection and robust optic nerve regeneration in mouse models of glaucoma and optic nerve injury.

## References

1. M. Malihi, E. R. Moura Filho, D. O. Hodge, A. J. Sit, Long-term trends in glaucoma- related blindness in Olmsted County, Minnesota. Ophthalmology 121, 134–141 (2014).

2. Y. Yokoyama et al., Characteristics of patients with primary open angle glaucoma and normal tension glaucoma at a university hospital: a cross-sectional retrospective study. BMC Res. Notes 8, 360 (2015).

3. D. M. O’Connor, N. M. Boulis, Gene therapy for neurodegenerative diseases. Trends Mol. Med. 21, 504–512 (2015).

4. A. M. Wilson, A. Di Polo, Gene therapy for retinal ganglion cell neuroprotection in glaucoma. Gene Ther. 19, 127–136 (2012).

5. R. Ren, Y. Li, Z. Liu, K. Liu, S. He, Long-term rescue of rat retinal ganglion cells and visual function by AAV-mediated BDNF expression after acute elevation of intraocular pressure. Invest. Ophthalmol. Vis. Sci. 53, 1003–1011 (2012).

6. S. G. Leaver et al., AAV-mediated expression of CNTF promotes long-term survival and regeneration of adult rat retinal ganglion cells. Gene Ther. 13, 1328–1341 (2006).

7. V. Pernet, A. Di Polo, Synergistic action of brain-derived neurotrophic factor and lens injury promotes retinal ganglion cell survival, but leads to optic nerve dystrophy in vivo. Brain 129, 1014–1026 (2006).

8. J. Dewitt et al., Constitutively active TrkB confers an aggressive transformed phenotype to a neural crest-derived cell line. Oncogene 33, 977–985 (2014).

9. E. C. Lerner et al., Ras CAAX peptidomimetic FTI-277 selectively blocks oncogenic Ras signaling by inducing cytoplasmic accumulation of inactive Ras-Raf complexes. J. Biol. Chem. 270, 26802–26806 (1995).

10. L. P. Wright, M. R. Philips, Thematic review series: lipid posttranslational modifications. CAAX modification and membrane targeting of Ras. J. Lipid Res. 47, 883–891 (2006).

11. T. Harada, C. Harada, L. F. Parada, Molecular regulation of visual system development: more than meets the eye. Genes Dev. 21, 367–378 (2007).

12. V. Gupta et al., BDNF impairment is associated with age-related changes in the inner retina and exacerbates experimental glaucoma. Biochim. Biophys. Acta 1842, 1567–1578 (2014).

13. A. Kimura, K. Namekata, X. Guo, C. Harada, T. Harada, Neuroprotection, growth factors and BDNF-TrkB signalling in retinal degeneration. Int. J. Mol. Sci. 17 (2016).

14. C. Harada et al., Glia- and neuron-specific functions of TrkB signalling during retinal degeneration and regeneration. Nat. Commun. 2, 189 (2011).

15. E. Sanz et al., Cell-type-specific isolation of ribosome-associated mRNA from complex tissues. Proc. Natl. Acad. Sci. U. S. A. 106, 13939–13944 (2009).

16. T. Harada et al., The potential role of glutamate transporters in the pathogenesis of normal tension glaucoma. J. Clin. Invest. 117, 1763–1770 (2007).

17. H. Sano et al., Differential effects of N-acetylcysteine on retinal degeneration in two mouse models of normal tension glaucoma. Cell Death Dis. 10, 75 (2019).

18. S. Honda et al., Survival of alpha and intrinsically photosensitive retinal ganglion cells in NMDA-induced neurotoxicity and a mouse model of normal tension glaucoma. Invest. Ophthalmol. Vis. Sci. 60, 3696–3707 (2019).

19. J. Zhang et al., Silicone oil-induced ocular hypertension and glaucomatous neurodegeneration in mouse. Elife 8 (2019).

20. X. Duan et al., Subtype-specific regeneration of retinal ganglion cells following axotomy: effects of osteopontin and mTOR signaling. Neuron 85, 1244–1256 (2015).

21. K. K. Park et al., Promoting axon regeneration in the adult CNS by modulation of the PTEN/mTOR pathway. Science 322, 963–966 (2008).

22. F. Bei et al., Restoration of visual function by enhancing conduction in regenerated axons. Cell 164, 219–232 (2016).

23. D. J. Creedon, E. M. Johnson, J. C. Lawrence, Mitogen-activated protein kinase- independent pathways mediate the effects of nerve growth factor and cAMP on neuronal survival. J. Biol. Chem. 271, 20713–20718 (1996).

24. Y. Zhou, V. Pernet, W. W. Hauswirth, A. Di Polo, Activation of the extracellular signal- regulated kinase 1/2 pathway by AAV gene transfer protects retinal ganglion cells in glaucoma. Mol. Ther. 12, 402–412 (2005).

25. Q. Wang et al., Optical control of ERK and AKT signaling promotes axon regeneration and functional recovery of PNS and CNS in Drosophila. Elife 9 (2020).

26. R. D. Almeida et al., Neuroprotection by BDNF against glutamate-induced apoptotic cell death is mediated by ERK and PI3-kinase pathways. Cell Death Differ. 12, 1329–1343 (2005).

27. M. Berry, Z. Ahmed, P. Morgan-Warren, D. Fulton, A. Logan, Prospects for mTOR- mediated functional repair after central nervous system trauma. Neurobiol. Dis. 85, 99–110 (2016).

28. T. Noro et al., Normal tension glaucoma-like degeneration of the visual system in aged marmosets. Sci. Rep. 9, 14852 (2019).

29. L. Cheng, P. Sapieha, P. Kittlerova, W. W. Hauswirth, A. Di Polo, TrkB gene transfer protects retinal ganglion cells from axotomy-induced death in vivo. J. Neurosci. 22, 3977–3986 (2002).

30. A. J. Weber, P. L. Kaufman, W. C. Hubbard, Morphology of single ganglion cells in the glaucomatous primate retina. Invest. Ophthalmol. Vis. Sci. 39, 2304–2320 (1998).

31. A. J. Weber, C. D. Harman, Structure-function relations of parasol cells in the normal and glaucomatous primate retina. Invest. Ophthalmol. Vis. Sci. 46, 3197–3207 (2005).

32. J. Agostinone et al., Insulin signalling promotes dendrite and synapse regeneration and restores circuit function after axonal injury. Brain 141, 1963–1980 (2018).

33. P. R. Williams, L. I. Benowitz, J. L. Goldberg, Z. He, Axon regeneration in the mammalian optic nerve. Annu. Rev. Vis. Sci. 6, 195–213 (2020).

34. S. de Lima et al., Full-length axon regeneration in the adult mouse optic nerve and partial recovery of simple visual behaviors. Proc. Natl. Acad. Sci. U. S. A. 109, 9149–9154 (2012).

35. J. H. Lim et al., Neural activity promotes long-distance, target-specific regeneration of adult retinal axons. Nat. Neurosci. 19, 1073–1084 (2016).

36. F. Sun et al., Sustained axon regeneration induced by co-deletion of PTEN and SOCS3. Nature 480, 372–375 (2011).

37. M. Leibinger et al., Boosting central nervous system axon regeneration by circumventing limitations of natural cytokine signaling. Mol. Ther. 24, 1712–1725 (2016).

38. R. Conceicao et al., Expression of developmentally important axon guidance cues in the adult optic chiasm. Invest. Ophthalmol. Vis. Sci. 60, 4727–4739 (2019).

39. X. W. Wang et al., Lin28 signaling supports mammalian PNS and CNS axon regeneration. Cell Rep. 24, 2540–2552 e2546 (2018).

40. C. Kole et al., Activating transcription factor 3 (ATF3) protects retinal ganglion cells and promotes functional preservation after optic nerve crush. Invest. Ophthalmol. Vis. Sci. 61, 31 (2020).

41. M. Leibinger, A. Andreadaki, H. Diekmann, D. Fischer, Neuronal STAT3 activation is essential for CNTF- and inflammatory stimulation-induced CNS axon regeneration. Cell Death Dis. 4, e805 (2013).

42. M. Leibinger et al., Transneuronal delivery of hyper-interleukin-6 enables functional recovery after severe spinal cord injury in mice. Nat. Commun. 12, 391 (2021).

43. T. Baden et al., The functional diversity of retinal ganglion cells in the mouse. Nature 529, 345–350 (2016).

44. B. A. Rheaume et al., Single cell transcriptome profiling of retinal ganglion cells identifies cellular subtypes. Nat. Commun. 9, 2759 (2018).

45. N. M. Tran et al., Single-cell profiles of retinal ganglion cells differing in resilience to injury reveal neuroprotective genes. Neuron 104, 1039–1055 (2019).

46. D. J. Liebl, W. Huang, W. Young, L. F. Parada, Regulation of Trk receptors following contusion of the rat spinal cord. Exp. Neurol. 167, 15–26 (2001).

47. Y. Zhu et al., Ablation of NF1 function in neurons induces abnormal development of cerebral cortex and reactive gliosis in the brain. Genes Dev. 15, 859–876 (2001).

48. K. Namekata et al., Dock3 induces axonal outgrowth by stimulating membrane recruitment of the WAVE complex. Proc. Natl. Acad. Sci. U. S. A. 107, 7586–7591 (2010).

49. B. M. Wice, J. I. Gordon, A strategy for isolation of cDNAs encoding proteins affecting human intestinal epithelial cell growth and differentiation: characterization of a novel gut-specific N-myristoylated annexin. J. Cell Biol. 116, 405–422 (1992).

50. H. Jiang et al., Protein lipidation: Occurrence, mechanisms, biological functions, and enabling technologies. Chem. Rev. 118, 919–988 (2018).

51. C. T. Rueden et al., ImageJ2: ImageJ for the next generation of scientific image data. BMC Bioinformatics 18, 529 (2017).

52. T. Udagawa et al., FUS regulates AMPA receptor function and FTLD/ALS-associated behaviour via GluA1 mRNA stabilization. Nat. Commun. 6, 7098 (2015).

53. S. Zolotukhin et al., Recombinant adeno-associated virus purification using novel methods improves infectious titer and yield. Gene Ther. 6, 973–985 (1999).

54. N. Itoh et al., Cell-specific and region-specific transcriptomics in the multiple sclerosis model: Focus on astrocytes. Proc. Natl. Acad. Sci. U. S. A. 115, E302–E309 (2018).

55. X. Guo et al., ASK1 signaling regulates phase-specific glial interactions during neuroinflammation. Proc. Natl. Acad. Sci. U. S. A. 119, e2103812119 (2022).

56. B. Langmead, S. L. Salzberg, Fast gapped-read alignment with Bowtie 2. Nat. Methods 9, 357–359 (2012).

57. D. Kim et al., TopHat2: accurate alignment of transcriptomes in the presence of insertions, deletions and gene fusions. Genome Biol. 14, R36 (2013).

58. S. Anders, P. T. Pyl, W. Huber, HTSeq--a Python framework to work with high-throughput sequencing data. Bioinformatics 31, 166–169 (2015).

59. S. Anders, W. Huber, Differential expression analysis for sequence count data. Genome Biol. 11, R106 (2010).

60. X. Mao, T. Cai, J. G. Olyarchuk, L. Wei, Automated genome annotation and pathway identification using the KEGG Orthology (KO) as a controlled vocabulary. Bioinformatics 21, 3787–3793 (2005).

61. Y. Azuchi et al., Role of neuritin in retinal ganglion cell death in adult mice following optic nerve injury. Sci. Rep. 8, 10132 (2018).

62. K. Namekata et al., DOCK8 is expressed in microglia, and it regulates microglial activity during neurodegeneration in murine disease models. J. Biol. Chem. 294, 13421–13433 (2019).

63. A. Erturk et al., Three-dimensional imaging of solvent-cleared organs using 3DISCO. Nat. Protoc. 7, 1983–1995 (2012).

64. X. Luo et al., Three-dimensional evaluation of retinal ganglion cell axon regeneration and pathfinding in whole mouse tissue after injury. Exp. Neurol. 247, 653–662 (2013).

